# Distinct interactions stabilize EGFR dimers and higher-order oligomers in cell membranes

**DOI:** 10.1101/2023.04.10.536273

**Authors:** Krishna C. Mudumbi, Eric A. Burns, David J. Schodt, Zaritza O. Petrova, Anatoly Kiyatkin, Lucy W. Kim, Emma M. Mangiacapre, Irais Ortiz-Caraveo, Hector Rivera Ortiz, Chun Hu, Kumar D. Ashtekar, Keith A. Lidke, Diane S. Lidke, Mark A. Lemmon

## Abstract

The epidermal growth factor receptor (EGFR) is a receptor tyrosine kinase (RTK) with important roles in many cellular processes as well as cancer and other diseases. EGF binding promotes EGFR dimerization and autophosphorylation through interactions that are well understood structurally. However, it is not clear how these dimers relate to higher-order EGFR oligomers detected at the cell surface. We used single-particle tracking (SPT) and Förster resonance energy transfer (FRET) imaging to examine how each domain within EGFR contributes to receptor dimerization and the rate of its diffusion in the cell membrane. We show that the EGFR extracellular region is sufficient to drive receptor dimerization, but that the EGF-induced EGFR slow-down seen by SPT requires formation of higher order oligomers, mediated in part by the intracellular tyrosine kinase domain – but only when in its active conformation. Our data thus provide important insight into higher-order EGFR interactions required for EGF signaling.

## Introduction

Receptor tyrosine kinases (RTKs) control many key cellular processes, and their dysfunction is associated with many human diseases^1^. Accordingly, efforts to understand transmembrane signaling mechanisms of RTKs have been considerable, with the epidermal growth factor receptor (EGFR) being among the most extensively studied^2–14^. Work over the past four decades has led to a prevailing model for EGFR signaling in which the unactivated receptor exists in a monomer-dimer equilibrium, shifting to the formation of active dimers upon binding of activating ligands^15, 16^. Structural studies explain how binding of ligands to the extracellular region (ECR) of EGFR (Fig. 1a) and its relatives induces conformational changes that promote homo- or heterodimerization^13, 17–19^. The resulting dimerization of the receptor allows the intracellular tyrosine kinase domains (TKDs) to form a characteristic asymmetric dimer^12^ in which one TKD allosterically activates its neighbor by inducing conformational changes in the TKD N-lobe^12^. Key tyrosines in the regulatory C-terminal tail (C-Tail) of EGFR (Fig. 1a) are then autophosphorylated in *trans*, and the resulting phosphotyrosines recruit downstream SH2 domain-containing signaling proteins to initiate an array of signaling cascades^1^.

**Fig. 1.**
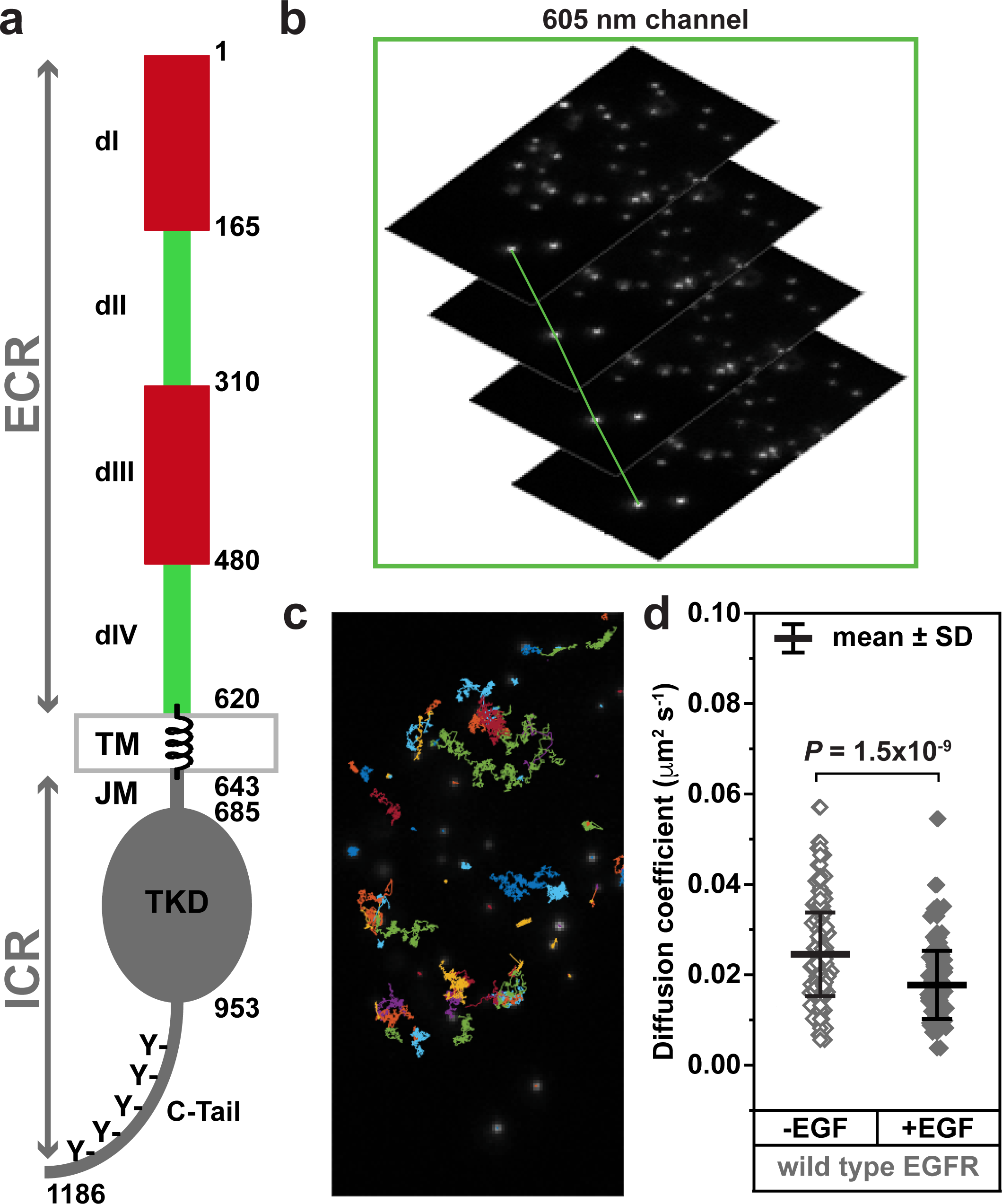
Single-particle tracking of EGFR in CHO cells. **a** Schematic diagram of epidermal growth factor receptor (EGFR) defining the major domains: extracellular region (ECR), intracellular region (ICR), transmembrane domain (TM), intracellular juxtamembrane domain (JM), tyrosine kinase domain (TKD), and the C-terminal tail (C-Tail). **b** Single-particle tracking (SPT) was performed using streptavidin-quantum dots (QD) conjugated to biotinylated HA antibody Fabs (QD-HA) or to biotinylated EGF (QD-EGF) as detailed in the Methods. Unless otherwise specified, cells were stimulated and tracked using QD-EGF at 200 pM. Time series were acquired at 46 Hz and individual receptors were tracked over time. **c** Representative 2D trajectories are shown for individual receptors diffusing on the cell membrane, from which diffusion coefficients (*D*) were calculated for each receptor and averaged over the cell (see Methods). **d** HA-tagged wild type EGFR expressed in CHO cells was tracked without ligand stimulation using QD-HA (gray open diamonds) or following stimulation with 200 pM QD-EGF (gray filled diamonds). The mean measured *D* value for each cell (n ≥ ∼50 tracks per cell) is plotted. Mean (± SD) across all cells are indicated (see Supplementary Table 1), and distributions are compared using Welch’s t-test (*P* = 1.5 × 10^-9^). All source data are provided as a Source data file.

Although structures of isolated monomers and dimers of the purified ECR and TKD from EGFR and its relatives are very well characterized^14, 15^, it remains a major challenge in the field to translate structural views of receptor dimerization into an understanding of how the receptor functions in cell membranes^20, 21^. Fluorescence- based imaging studies in cells have suggested that EGFR forms ligand-independent oligomers^20, 22, 23^. There is also evidence for ligand-induced oligomers beyond dimers, depending on the cellular context^20, 22–26^. Although there is good agreement on the structural basis for simple receptor dimerization, models for the various proposed higher order oligomers vary widely in structural detail^21, 24, 25^. Moreover, it is not clear what role each individual domain within EGFR (Fig. 1a) plays in stabilizing dimers or oligomers of the full-length receptor.

To address some of these questions, we used single-particle tracking (SPT), Förster resonance energy transfer-fluorescence lifetime imaging microscopy (FRET-FLIM), super-resolution microscopy, and cellular signaling studies to interrogate the relative roles of the different parts of EGFR in both EGFR dimerization and higher order oligomerization. SPT and FRET studies have been used extensively to monitor receptor dynamics and interactions in cell membranes^2, 27–29^, and have the advantage of being able to monitor variants of EGFR with well-defined structural changes directly in the complex milieu of the cell membrane^2, 27–31^. We find that the reduction in EGFR mobility typically seen upon ligand binding requires receptor dimerization. An intact ligand-bound ECR is sufficient for EGFR dimerization, but dimerization of the receptor alone is not sufficient for the observed slow-down in SPT studies. In the context of ligand-induced EGFR dimers, we find that interactions involving the intracellular JM and TKD domains are crucial for the slow-down of EGFR, which we also find requires the TKD to adopt an active-like conformation. Our data therefore reinforce recent suggestions that EGFR oligomers (dimers of EGF-induced dimers) are important for signaling. They also provide a valuable platform for resolving disagreements between structural models for these receptor oligomers and thus to identify potential new approaches for EGFR inhibition.

## Results

Although detailed conclusions drawn from SPT studies of EGFR have differed, depending in part on the question asked, they all find that EGF binding reduces receptor mobility in the plasma membrane^2, 20, 27–31^. This effect is measured as a change in the lateral diffusion coefficient, *D*, and is associated with receptor activation^20^. The observed reduction in mobility has variously been argued to reflect receptor dimerization itself^2^, a more complex function of receptor dimerization/oligomerization, recruitment of downstream signaling molecules, in addition to changes in the membrane and/or cytoskeletal environment^2, 7, 32^. All studies agree, however, that receptor dimerization of the sort that has been described structurally^14, 15^ is necessary for the observed reduction in EGFR mobility upon ligand binding, and mutations that impair the extracellular dimerization arm, for example, diminish the observed slow-down^2, 31^.

We used quantum dot (QD)-conjugated HA tag antibody Fab fragments to label N-terminally HA-tagged EGFR (HA-EGFR) expressed at the surface of transfected CHO cells, which lack endogenous EGFR^33, 34^. Using total internal reflection microscopy (TIRFM), we imaged the ligand-free receptor at the cell surface for 1 min (Fig. 1b,c), imaging for no more than a total of 10 min per condition per dish. Under these conditions, SPT analysis of unliganded EGFR yielded a mean diffusion coefficient (*D*) of 0.025 ± 0.009 μm^2^s^-1^ (Fig. 1d and Supplementary Table 1). When the same experiment was performed using QD-labeled EGF^31, 35^ (QD-EGF) to label the receptor, however, the mean value of *D* for EGF-bound EGFR fell to 0.018 ± 0.008 μm^2^s^-1^ (*P* = 1.5 × 10^-9^), in line with previous studies of the ability of EGF to reduce receptor mobility^2, 7, 30–32^. A similar change was seen when receptor labeled with QD-conjugated Fab fragments was instead compared in the absence and presence of excess unlabeled EGF (Supplementary Table 2), with a fall in *D* in this case from 0.022 ± 0.012 μm^2^s^-1^ to 0.012 ± 0.008 μm^2^s^-1^ (*P* = 5.5 × 10^-6^). Cells were sorted before use in all experiments to achieve approximately constant (and physiological) levels of EGFR expression (∼5,000- 10,000 EGFRs/cell). We also performed additional controls to determine whether expression level or receptor location (basolateral versus apical) influenced the observed EGF-induced receptor slow-down (Supplementary Figure 1a,b). No significant effects were observed.

### EGFR mobility reflects extracellular region dimerization strength

It was previously shown that abolishing EGFR dimerization by mutating the key dimerization arm in the extracellular region (ECR) of the receptor prevents EGF from reducing receptor diffusion at the cell surface^31^. Similarly, treatment with a ligand that induces formation of only weak EGFR dimers (e.g. epiregulin, or EREG) fails to strongly slow down the wild type EGFR^30^. We recently showed that key EGFR ECR mutations found in glioblastoma patients (R84K and A265V) strengthen the ECR dimer induced by EREG so that it resembles an EGF-induced dimer^18^. These mutations provide an opportunity to ask how strengthening the ECR dimer in intact EGFR within cells affects its diffusion. When EGFR mobility is monitored with QD-conjugated HA-Fab, adding saturating amounts of unlabeled EREG (1 μM) leads to only a small reduction in *D* that is barely significant (*P* = 0.044; Supplementary Table 2), whereas saturating levels of unlabeled EGF (50 nM) induce the expected near-halving of the *D* value (Fig. 2 and Supplementary Table 2). By contrast, *D* values for the R84K- and A265V-mutated EGFRs are reduced by 43% (*P* = 6.0 × 10^-5^) and 31% (*P* = 1.4 × 10^-4^) respectively (Fig. 2; Supplementary Table 2) by EREG – similar to the effects seen with EGF for the wild type or GBM-mutated receptor. Beyond confirming that dimerization is necessary for the reduced rates of diffusion seen upon ligand binding to EGFR, these data reveal a clear correlation between diffusion coefficients measured for intact EGFR in live cells and the strength of dimerization of the isolated ECR as measured using biophysical methods in solution^18, 30^.

**Fig. 2.**
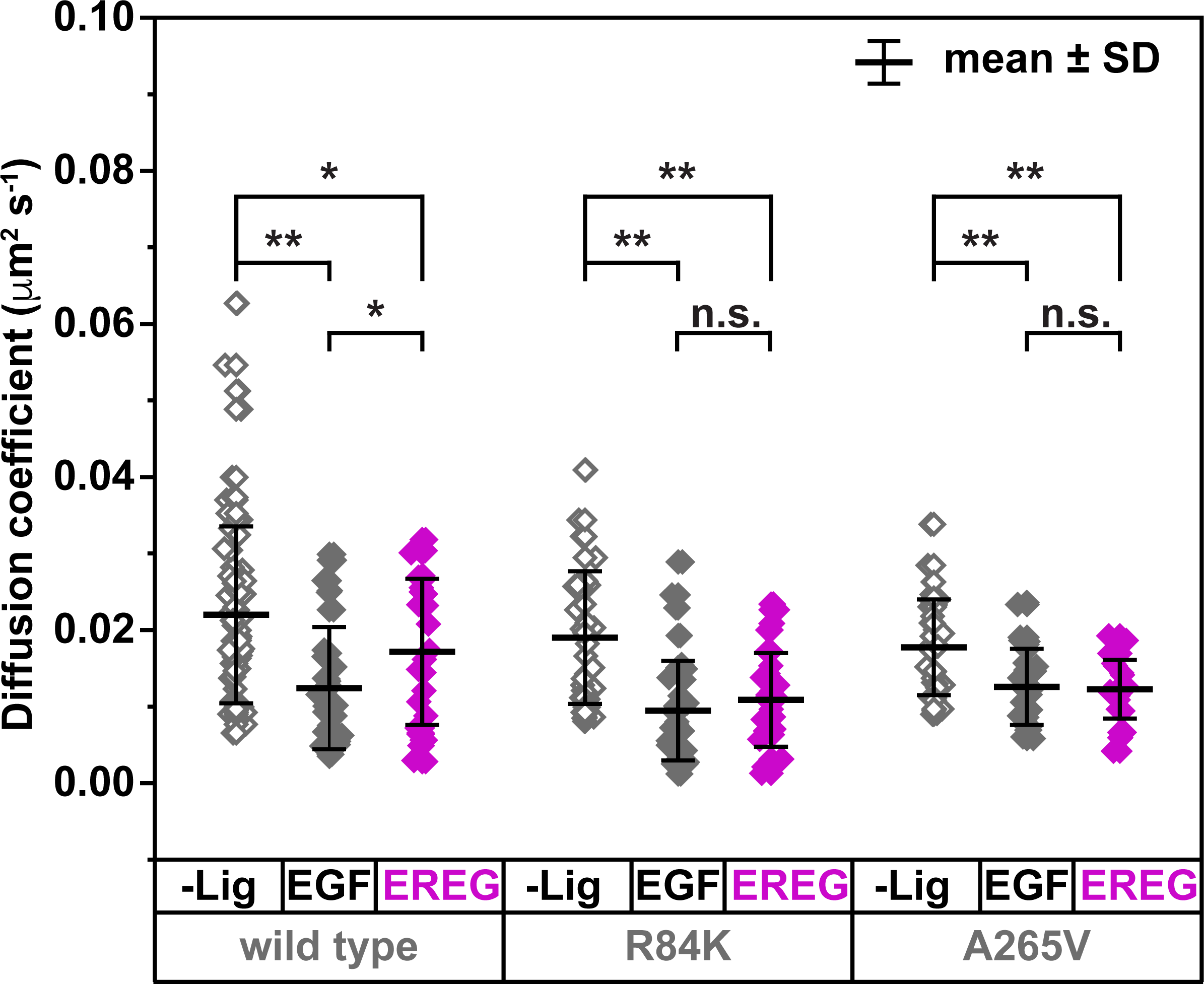
Altered ECR dimerization strength directly affects EGFR diffusion. SPT of wild type EGFR and variants harboring extracellular mutations (R84K and A265V) found in glioblastoma multiforme (GBM) was performed in the absence of ligand (gray open diamonds), after stimulation with 50 nM EGF (gray filled diamonds), and following stimulation with the low affinity ligand epiregulin EREG at 1 μM (magenta filled diamonds). Unlabeled (dark) ligands were used for these experiments, and all receptors were tracked by labeling with QD-HA. Wild type EGFR shows a significantly smaller reduction in mean *D* value with EREG (*n* = 26) than with EGF (*n* = 33; *P* = 0.047), consistent with weakened dimerization^30^, whereas both the R84K and A265V variants are affected similarly by both ligands (*P* values of 0.4 and 0.8 respectively, with *n* ≥ 27 and 21, see Supplementary Table 2). Unpaired two-sided Welch’s t-tests were used to calculate *P* values (\**P* < 0.05; \*\**P* < 1×10^-3^; \*\*\**P* < 1×10^-6^), which are all listed in Supplementary Table 2, and n.s. indicates no significant difference. All source data are provided as a Source data file.

### TM domain dimerization motifs do not influence EGFR diffusion

There has been substantial discussion in the literature about the role of the transmembrane (TM) domain of EGFR in receptor dimerization and signal transmission. Early studies indicated that the TM domain is highly tolerant to substitutions without detrimental effects on signaling^36, 37^. Later studies identified three Sm-x-x-x-Sm TM domain dimerization motifs^38^ (Fig. 3a) in EGFR (where Sm represents any amino acid with a small side-chain, allowing close crossing of TM helices^39^), and found that they contribute to dimerization of the isolated TM domain when studied in bacterial membranes^40^. Although never visualized structurally in any receptor context, it was later argued that different interactions between these Sm-x-x-x-Sm motifs within the TM domain promote dimerization in active and inactive EGFR dimers^41^, with effects on signaling when mutated. Other signaling studies^42, 43^ argue that none of the TM dimerization motifs are required for signaling – consistent with earlier views. Given the different viewpoints in the field, we were interested in determining whether the dimerization motifs in the EGFR TM domain are necessary for the ligand-induced EGFR mobility reductions seen in SPT studies. As shown in Fig. 3b, complete removal of the Sm-x-x-x-Sm TM domain dimerization motifs from the EGFR TM domain (in TM3X, where the small residues are replaced with larger valines: Fig. 3a) did not prevent ligand-induced changes in diffusion – with the expected ∼2-fold reduction being clearly seen upon EGF addition (Supplementary Table 3). Although it seems highly likely that TM domain interactions do contribute to self-association of EGFR^40, 42^ and other single-TM receptors^44^, our data and those of Bartzoka *et al*.^42^ argue that they are not required for ligand-induced EGFR dimerization or signaling as assessed by EGF-induced EGFR autophosphorylation (Fig. 3c), suggesting that they play a more passive role^43^ in the wild type receptor.

**Fig. 3.**
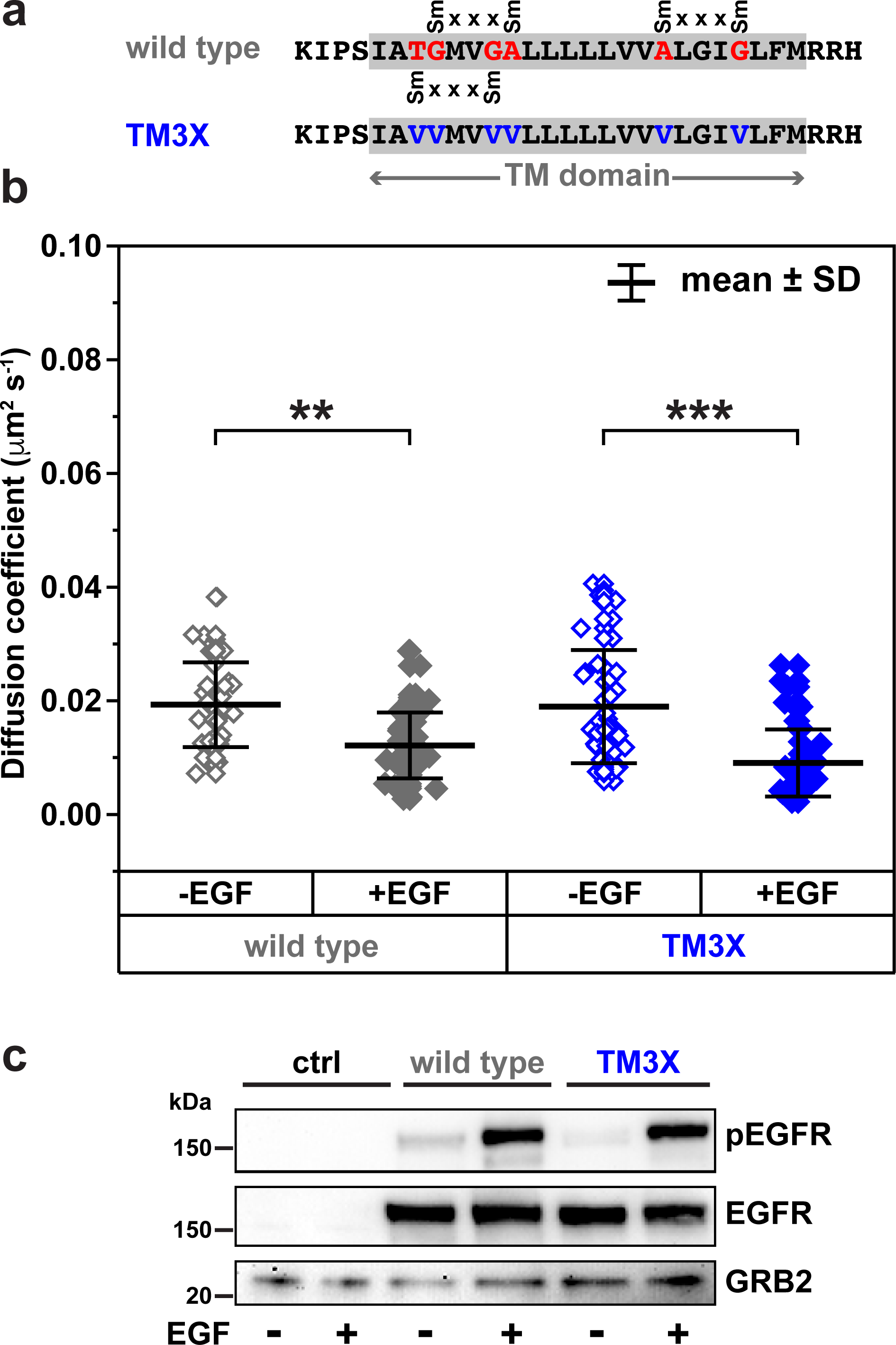
TM domain dimerization motifs do not influence EGFR diffusion. **a** Schematic of mutations introduced to disrupt the Sm-x-x-x-Sm dimerization motifs^38^ in the EGFR TM domain. There are three Sm-x-x-x-Sm motifs, where Sm is any small amino acid, in EGFR (T-g-m-v-G; G-m-v-g-A; and A-l-g-i-G), with the small residues colored red. All Sm residues were mutated to valine in TM3X. **b** SPT data for wild type EGFR (gray) and a variant harboring TM mutations that block TM domain dimerization^40^ (TM3X: blue). HA-tagged receptors were labeled and tracked with QD-HA in the absence of ligand (open diamonds), and using 200 pM QD-EGF to investigate the effect of ligand binding (filled diamonds). EGF induces a very similar and significant slow-down for both wild type EGFR (*n* = 40 without ligand, 42 with ligand; *P* = 6.7 × 10^-6^: see Supplementary Table 3) and the TM3X variant (*n* = 45 without ligand, 46 with ligand; *P* = 1.9 × 10^-7^). Unpaired two-sided Welch’s t-tests were used to calculate *P* values (\*\**P* < 1×10^-3^; \*\*\**P* < 1×10^-6^). **c** pEGFR immunoblots of wild type and TM3X EGFR activated with 16 nM EGF for 5 min and probed with anti-pY1173 (upper), anti-EGFR (middle) and anti-GRB2 (lower) as loading control. Representative of three biological repeats. All source data and uncropped gels (including total EGFR and GRB2 and tubulin loading controls) are provided in a Source data file.

### The intracellular region (ICR) is required for altered receptor mobility, but not for dimerization

We were next interested in understanding the relative contributions of extra- and intra- cellular interactions to EGFR dimerization in cell membranes, about which there has been some debate in the literature^6, 15, 41, 45–47^. We previously showed that the isolated ECR of EGFR dimerizes with a dissociation constant (*K*_d_) of 1-3 μM when bound to EGF or TGFα^46, 47^. An interaction with this strength should be sufficient to allow the ECR alone to drive dimerization of most EGF-bound EGFR in typical cells that have several thousand receptors, where local EGFR concentration is estimated at 1-10 μM^48^ – even ignoring pre-orientation effects in the membrane (which will further enhance dimerization^49, 50^). Contrary to this view, however, others have argued instead that a strong intrinsic dimerization ability of the cytoplasmic module or ICR of EGFR drives receptor dimerization^6, 41, 45^ – and that this tendency of the ICR to dimerize is suppressed by the ECR until extracellular ligand binding relieves inhibitory contacts. In support of this argument, Jura *et al*.^6^ estimated that an EGFR fragment containing just the juxtamembrane (JM) region and TKD of the receptor dimerizes with a *K*_d_ value of ∼0.2 μM by monitoring kinase activity as a function of concentration – arguing that this intracellular module dimerizes more strongly than the EGF-bound ECR. We were therefore interested to use SPT to analyze an EGFR variant that lacks its cytoplasmic module (ΔICR; truncated after residue 670 in mature EGFR numbering). As shown in Fig. 4a, EGFR(ΔICR) differed from intact EGFR by displaying no significant EGF-induced reduction in diffusion (*P* = 0.21). Measured *D* values for EGFR(ΔICR) were 0.026 ± 0.013 μm^2^s^-1^ in the absence of ligand, and 0.028 ± 0.013 μm^2^s^-1^ with 200 pM QD-EGF (Supplementary Table 3).

**Fig. 4.**
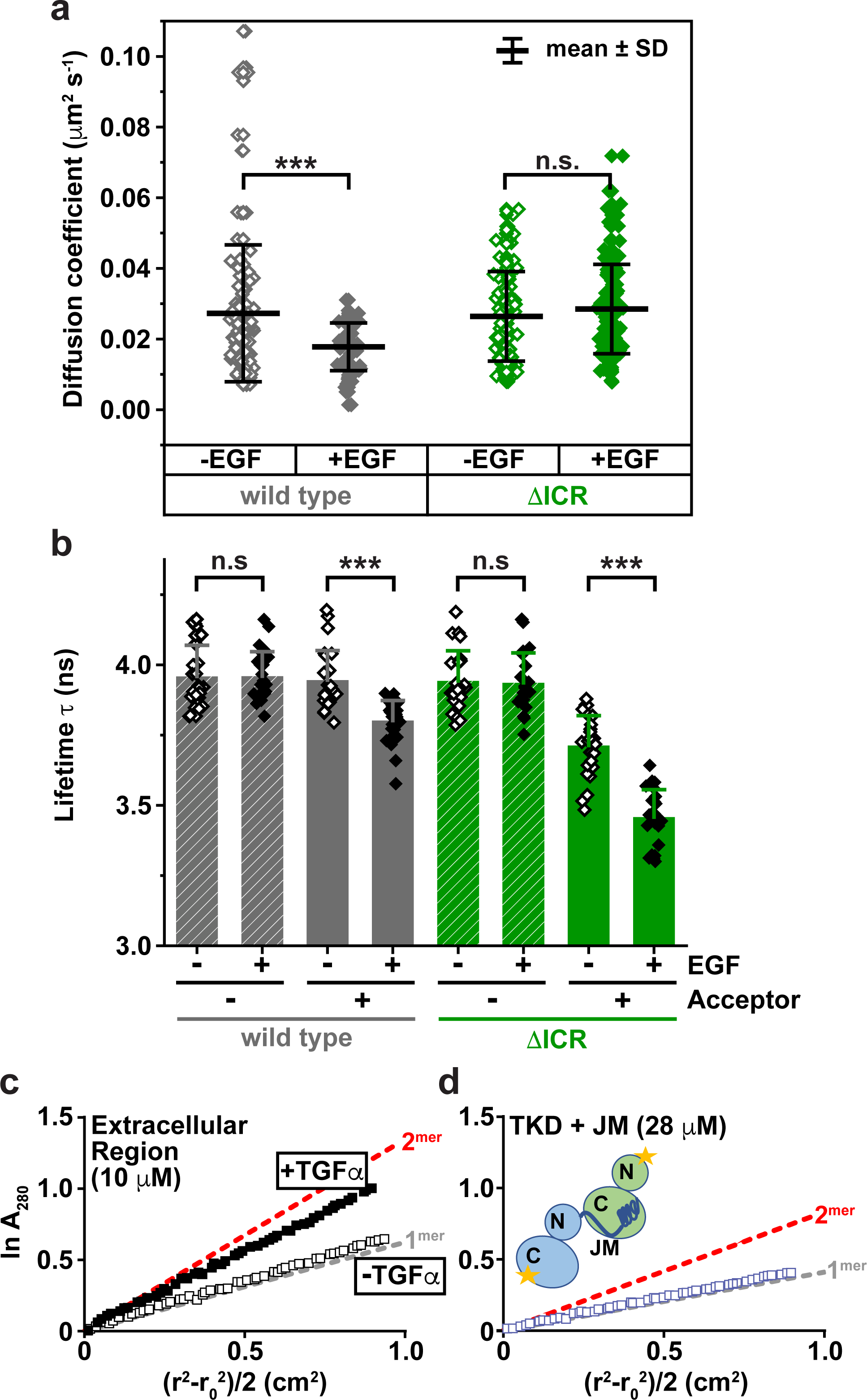
TKD deletion prevents ligand-induced slow-down but not dimerization. **a** SPT analysis of wild type EGFR (gray) and a truncation variant (ΔICR: green) in which the entire intracellular region has been removed (residues 671-1186 in mature EGFR; 695-1210 in pro-EGFR). Wild type EGFR shows a significant reduction in mean *D* value upon QD-EGF (200 pM) binding (*P* = 9.2 × 10^-6^; *n* = 124 without ligand, 56 with ligand: see Supplementary Table 3), but this is lost when the ICR is deleted (*P* = 0.21; *n* = 116 without ligand, 121 with ligand). Unpaired two-sided Welch’s t-tests were used to calculate *P* values (\*\*\**P* < 1×10^-6^; n.s. indicates no significant difference). **b** FRET-FLIM imaging analysis of HaloTagged wild type or ΔICR EGFR expressed in CHO cells as described in Methods. Imaging experiments were performed both after labeling with the donor fluorophore (JF-549; 20 nM) only and after labeling with both donor and acceptor (JF-646) fluorophores at equal concentrations (20 nM). Imaging was performed with (+) and without (-) addition of 50 nM EGF (unlabeled) to ensure saturation. Unpaired two-sided Welch’s t-tests were used to calculate *P* values (\*\*\**P* < 1×10^-6^), with n.s. indicating no significant difference. Donor fluorescence lifetimes were reduced significantly (with donor and acceptor) upon EGF addition for both wild type EGFR (*P* = 8.1×10^-7^; *n* = 25 without ligand, 29 with ligand) and ΔICR (*P* = 1.3×10^-11^; *n* = 26 without ligand, 24 with ligand). **c** Analytical ultracentrifugation (AUC) sedimentation-equilibrium (SedEq) analysis of soluble ECR at 10 μM without ligand (open black boxes), and with addition of a 1.2-fold excess of TGFα (filled black boxes) – chosen because it absorbs much less than EGF at 280 nm, facilitating data interpretation^46^. Data are presented in logarithmic form, where a single species gives a straight line with gradient proportional to molecular weight. The superimposed gray dashed line corresponds to expected data for the monomeric ECR, and red for the ECR dimer. As in previous studies^46^, TGFα induces almost complete ECR dimerization, consistent with a *K*_d_ value of 1-3 μM. Representative of three biological repeats. **d** SedEQ-AUC analysis of EGFR ICR dimerization. Two purified fragments of the EGFR ICR were mixed at 1:1 ratio, with each present at 14 μM. One was a TKD construct with the complete JM region plus a V924R mutation that allows it to interact with another TKD only through its N-lobe (see Fig. 5a). The other lacks the JM, but has an I682Q mutation that allows interaction with another TKD only through the C-lobe. Mixing I682Q and V924R mutated receptors allows discrete formation of the asymmetric dimer^12^, which we wish to measure. Gray and red lines represent monomer and dimer as in **c**, and the protein mixture runs at monomer molecular weight, arguing the *K*_d_ for this interaction is >100 μM. Representative of three biological repeats. All source data are provided as a Source data file.

Given the literature arguments summarized above, the failure of EGF to reduce mobility of EGFR(ΔICR) might be taken to suggest that this construct fails to dimerize upon EGF binding. Since this result would be very surprising given the ECR dimerization strength and the receptor expression levels used here (∼5.000-10,000 receptors/cell), we also monitored receptor dimerization using an orthogonal approach. We used Förster resonance energy transfer-fluorescence lifetime imaging microscopy (FRET-FLIM) with N-terminally HaloTagged receptor variants labeled with Janelia Fluor® HaloTag® ligands: JF549 as FRET donor and JF646 as FRET acceptor. As shown in Fig. 4b, donor fluorescence lifetime (τ) was significantly reduced (in the presence of acceptor) when unlabeled EGF (50 nM) was added to either EGFR(ΔICR) (*P* = 1.3 × 10^-11^) or intact EGFR (*P* = 8.1 × 10^-7^) – arguing that EGFR(ΔICR) does in fact dimerize (at least) just as robustly as the wild type receptor upon EGF binding. These findings argue both that ICR interactions are not required for EGFR dimerization and that (ECR-mediated) dimerization of the receptor is not sufficient on its own for the observed EGF-induced reduction in EGFR diffusion in cells. It is also noteworthy that the ΔICR construct exhibits a change in fluorescence lifetime even before addition of ligand, suggesting that the kinase domain and/or the C-Tail may have an autoinhibitory effect that reduces dimerization in the absence of ligand.

To further compare quantitative contributions of the ECR and ICR to EGFR dimerization, we used sedimentation equilibrium analytical ultracentrifugation (SedEq- AUC) to assess dimerization strengths. As in our previously published studies^46, 47, 51^, saturating the isolated ECR of the receptor (at 10 μM) with ligand causes it to dimerize almost completely (Fig. 4c), consistent with the published *K*_d_ value of ∼1 μM^46^. By contrast, formation of the asymmetric ICR dimer could not be detected at all using a JM- plus-TKD construct even at 28 μM (Fig. 4c). These experiments utilized mutated TKD variants designed to measure formation of discrete asymmetric dimers only, rather than aggregates (see legend to Fig. 4c), and suggest that the *K*_d_ value for dimerization of the JM-plus-TKD region of EGFR exceeds 100 μM. These data therefore argue that the ECR dominates in driving ligand-induced EGFR dimerization.

### Mutations in the JM latch or asymmetric dimer interface prevent EGF-induced slow-down

Although the above experiments argue that EGFR dimerization is primarily driven by the ECR, they also show that the ICR is required for the observed ligand-induced reduction in receptor diffusion in cells. Fig. 5a gives a structural view of the asymmetric dimer formed between two ICRs, with interactions mediated by the JM region and N-lobe of the receiver TKD (gray/cyan) and the C-lobe of the activator TKD (yellow; interface in purple)^6, 9, 12^. We next asked whether the specific interactions seen here are important for the EGF-induced reduction in EGFR diffusion. We first replaced (with alanine) four key residues in the JM region of the ‘receiver’ ICR (cyan in Fig. 5a) that participate in stabilizing the asymmetric dimer (L664/V665/L668/S671). The resulting variant (JM4X) no longer showed a significant reduction in diffusion coefficient upon EGF binding (Fig. 5b; Supplementary Table 3). Importantly, this EGFR variant could still be activated to a limited extent, as assessed both by autophosphorylation assays and analysis of ERK signaling (Fig. 5c), although activation was substantially suppressed compared with the wild type receptor. These data suggest that weakening JM-mediated ICR interactions allows the receptor to retain at least some key signaling events, but prevents interactions that normally lead to reduced receptor diffusion. Interestingly, EGF-activated JM4X may resemble weakly dimeric EREG-activated wild type EGFR in this respect, which retains ERK signaling (although with different kinetics^30^) but does not have a substantially reduced diffusion coefficient (Fig. 2). He and Hristova^52^ previously showed that complete replacement of the JM region (removing all JM-mediated interactions in Fig, 5a) abolishes receptor activation altogether, without preventing EGF-induced dimerization of the mutated receptor – consistent with our findings.

**Fig. 5.**
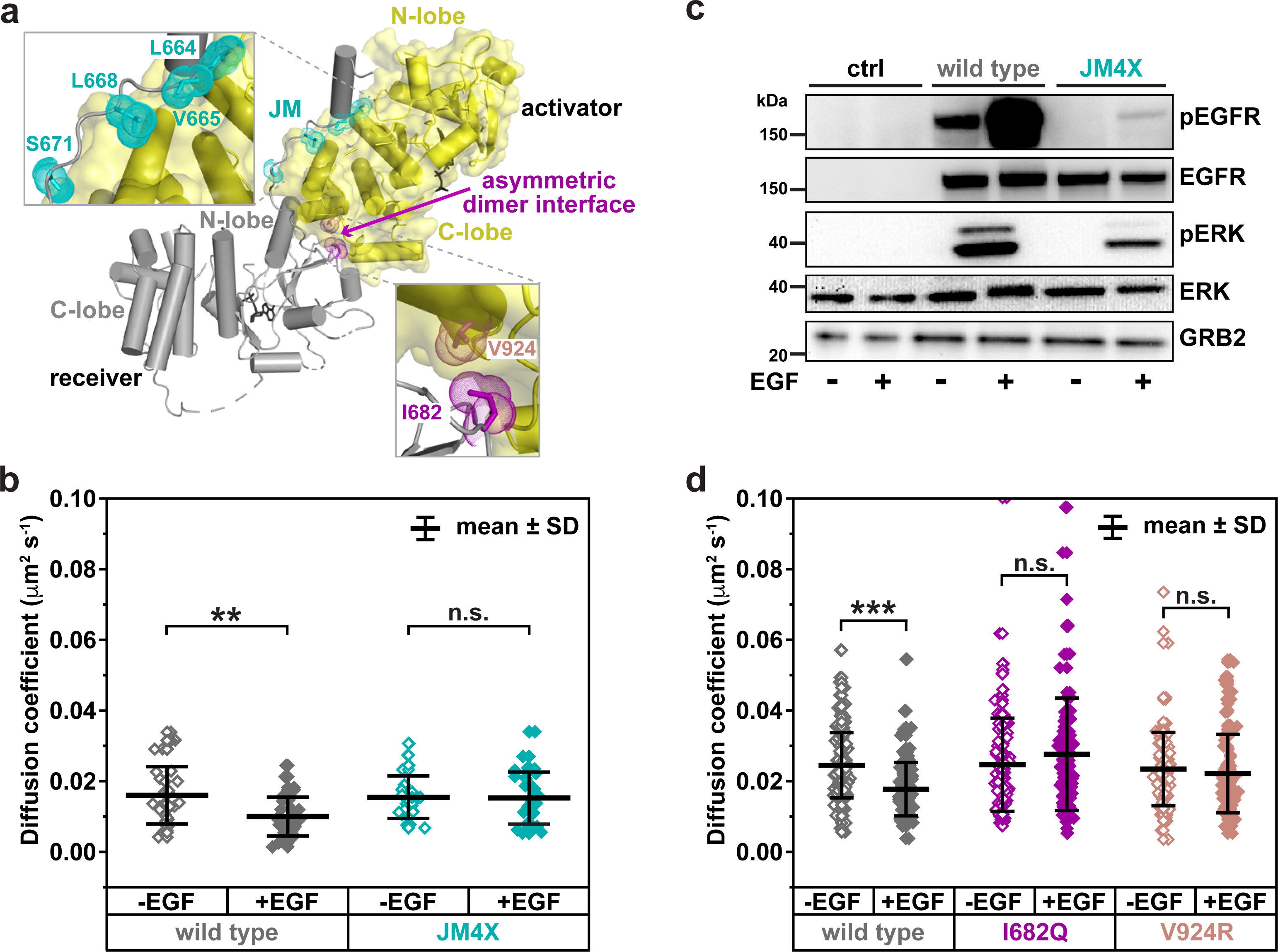
The JM latch and asymmetric kinase dimer are required for reduced EGFR diffusion. **a** Structural view of asymmetric dimer between two EGFR TKDs from PDB ID: 3GOP^9^. The C-lobe of the activator kinase (yellow) interacts with the N-lobe of the receiver kinase, leading to its allosteric activation^12^. The asymmetric dimer interface between receiver and activator is augmented by the JM ‘latch’^6,9^, with key interacting residues colored cyan (see left insert). The JM4X mutation includes replacing L664, V665, L668, and S671 all with alanine to disrupt JM interactions. Mutations to disrupt the asymmetric dimer interface are I682Q in the receiver N-lobe (purple) and V924R in the activator C-lobe (dark pink). **b** SPT analysis of wild type EGFR (gray) and the JM4X variant (cyan). QD-EGF (200 pM) induces a significant slow-down for wild type (*P* = 9.3 × 10^-4^; *n* = 32 without ligand, 34 with ligand), but this is lost in the JM4X variant (*P* = 0.91; *n* = 24 without ligand, 30 with ligand). Unpaired two-sided Welch’s t-tests were used to calculate *P* values (\*\**P* < 1×10^-3^; n.s. indicates no significant difference), listed in Supplementary Table 3. **c** pEGFR and pERK immunoblots of wild type and JM4X EGFR activated with 16 nM EGF for 5 min and probed with anti-pY1173, anti-EGFR, anti-pERK, anti-ERK, and anti- GRB2 (lower) as loading control. Source data and uncropped gels are provided in the Source data file. Representative of three biological repeats. The JM4X still shows a small degree of signaling at pEGFR and pERK levels, but it is greatly reduced compared with wild type. **d** SPT analysis of wild type EGFR (gray), the I682Q N-lobe-mutated variant (purple) and the V924R C-lobe mutated variant (dark pink). QD-EGF (200 pM) induces a significant slow-down for wild type in these experiments (*P* = 1.9 × 10^-9^; *n* = 126 without ligand, 118 with ligand), but this is lost in both the I682Q variant (*P* = 0.13; *n* = 116 without ligand, 116 with ligand) and the V924R variant (*P* = 0.36; *n* = 108 without ligand, 124 with ligand). Unpaired two-sided Welch’s t-tests were used to calculate *P* values (\*\*\**P* < 1×10^-6^; n.s. indicates no significant difference), listed in Supplementary Table 1. All source data are provided as a Source data file.

We also investigated the effects of key mutations in the N-lobe of the receiver TKD (I682Q) and the C-lobe of the ‘activator’ TKD (V924R) that disrupt the asymmetric dimer interface shown in Fig. 5a, and have been shown to largely abolish EGFR activation^12^. As shown in Fig. 5d and Supplementary Table 1, either of these mutations abolished the EGF-induced slow-down of EGFR. In the absence of EGF, mean *D* values were 0.025 ± 0.013 μm^2^s^-1^ (I682Q) and 0.023 ± 0.010 μm^2^s^-1^ (V924R). When analyzed with QD-EGF, *D* values were 0.028 ± 0.016 μm^2^s^-1^ (I682Q) and 0.022 ± 0.011 μm^2^s^-1^ (V924R). Taken together, these data argue that, although intracellular interactions mediated by the JM region or TKD are not required for dimerization, weakening (or abolishing) them prevents EGFR from engaging in the processes that lead to its reduced diffusion upon EGF binding.

### Reduced EGFR diffusion depends on TKD conformation but not activity

Having established that interactions involving the TKD are important for the EGF-induced reduction in EGFR diffusion, we next asked whether kinase activity is also required. Indeed, one possible explanation for the EGF-induced slow-down would be that recruitment of downstream signaling molecules to autophosphorylated tyrosines in EGFR lead to a large increase in size of the complex and a resulting reduction of mobility^7^. We used two approaches to investigate this question. In the first, we generated a ‘kinase dead’ EGFR (Kin^-^) by mutating the aspartate of the HRD motif in the TKD (D813 – in mature EGFR numbering) to asparagine. This mutation removes the catalytic base required for catalysis of phosphotransfer with minimal structural alteration^53^. In another approach, we used EGFR-specific tyrosine kinase inhibitors (TKIs) that bind to the ATP-binding site. As shown in Fig. 6 and Supplementary Table 4, Kin^-^ EGFR still showed a significant EGF-induced reduction in diffusion coefficient (*P* = 1.2 × 10^-4^), from 0.021 ± 0.006 μm^2^s^-1^ to 0.016 ± 0.006 μm^2^s^-1^. This result argues that the EGF-induced slow-down of EGFR does not require the receptor’s kinase activity, consistent with previously published observations by Low-Nam *et al*.^7^ and by Huang *et al*^24^.

**Fig. 6.**
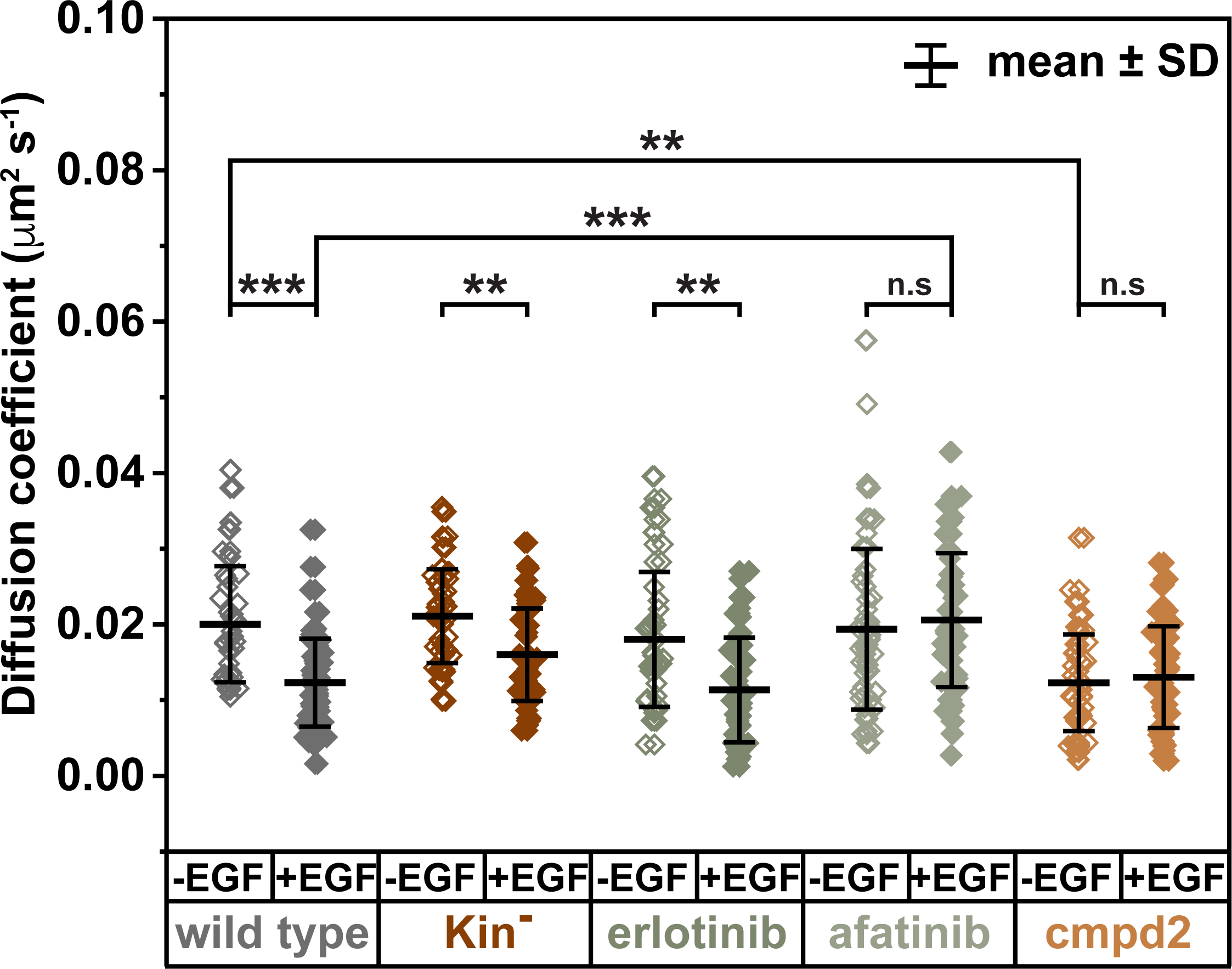
TKD conformation, not activity, influences EGFR diffusion. SPT analysis with and without QD-EGF (200 pM) of wild type EGFR (gray), ‘kinase dead’ (D813N; Kin^-^, brown), as well as wild type EGFR treated with inhibitors (each at 10 μM) that have different conformation state preferences. QD-EGF induces a significant slow-down for wild type (gray: *P* = 1.0 × 10^-6^; *n* = 39 without ligand, 63 with ligand), for Kin^-^ (brown: *P* = 1.2 × 10^-4^; *n* = 48 without ligand, 46 with ligand), and wild type with erlotinib (medium gray: *P* = 3.0 × 10^-4^; *n* = 39 without ligand, 46 with ligand). This effect is lost with afatinib (light gray: *P* = 0.52; *n* = 53 without ligand, 52 with ligand), which stabilizes a conformation inconsistent with formation of the asymmetric TKD dimer^56^. With compound 2 (Cmpd2), by contrast, inhibitor addition alone is sufficient to cause the slow-down (gold: *n* = 43 without ligand, 50 with ligand), consistent with the ability of this compound to stabilize the active conformation of the TKD^57^. Unpaired two-sided Welch’s t-tests were used to calculate *P* values (\*\**P* < 1×10^-3^; \*\*\**P* < 1×10^-6^), and n.s. indicates no significant difference. Data are summarized in Supplementary Table 4. All source data are provided as a Source data file.

To investigate further the effect of TKD conformation on EGFR diffusion while also inhibiting kinase activity, we used three different inhibitors known to have distinct effects on TKD conformation^54^.

#### Erlotinib

We first assessed the effect of erlotinib (added at 10 μM), which has no preference for one EGFR TKD conformation over another^55^. Although erlotinib was first visualized bound to the EGFR TKD in its active conformation – when soaked into pre-formed crystals of the asymmetric TKD dimer^11^ – co-crystallization and computational modeling studies^55^ revealed that it binds equally well to the inactive conformation. Saturating levels of erlotinib do not significantly alter receptor mobility, and the EGF-induced slow- down is retained (Fig. 6 and Supplementary Table 4), with *D* falling by 37% upon EGF addition (*P* = 3 × 10^-4^).

#### Afatinib

In stark contrast to erlotinib, the second-generation covalent EGFR inhibitor afatinib (added at 10 μM) abolished the EGF-induced slow-down of EGFR (Fig. 6 and Supplementary Table 4), with *D* remaining at 0.021 ± 0.009 μm^2^s^-1^ (*P* = 0.5) after EGF addition. When co-crystallized with EGFR TKD (in PDB entry 4G5P^56^) rather than being soaked into pre-existing active dimer crystals, afatinib binding prevents the TKD from making the normal asymmetric TKD dimer contacts, suggesting that afatinib binding disrupts the active conformation. Distorting the active conformation of the TKD with afatinib may therefore prevent the EGF-induced reduction in EGFR mobility.

#### Compound 2

Consistent with the suggestion that the active TKD conformation is required for EGFR slow-down, we found that a compound known to robustly stabilize the active EGFR TKD conformation – called compound 2 (also at 10 μM)^57^ – reduces EGFR diffusion (to a *D* value of 0.012 ± 0.006 μm^2^s^-1^) even in the absence of ligand (Fig. 6 and Supplementary Table 4). Adding EGF causes no significant further reduction (*D* = 0.013 ± 0.007 μm^2^s^-1^; *P* = 0.6). Compound 2 was identified in a screen for small molecules that inhibit signaling by ErbB3/ErbB2 heterodimers^57^. We showed crystallographically that compound 2 stabilizes the active conformation of EGFR^57^ and thus presumably the asymmetric TKD dimer. This result resembles our previous finding that activating TKD mutations seen in lung cancer also slow down the receptor^31^, presumably also by stabilizing the active TKD conformation. These data are also consistent with the increase in EGFR homodimerization reported upon gefitinib binding^58^ – although the strong selectivity of compound 2 for binding the active EGFR TKD conformation^57^ makes the effect more complete in this case.

Taken together, the observations in Figs. 5 and 6 argue that the EGF-induced EGFR slow-down requires formation of the intracellular asymmetric active-like TKD dimer in the context of an ECR-driven receptor dimer. This can occur in the absence of tyrosine kinase activity (e.g. with the Kin^-^ variant or with addition of erlotinib), and can be prevented by mutations in the asymmetric TKD dimer interface or by stabilization of incompatible TKD conformations. On the other hand, the slow-down can be promoted even in the absence of EGF by inhibitors that stabilize the active TKD conformation (compound 2) or by mutations that promote dimerization (L858R or exon 19 deletions^31^).

### The influence of autophosphorylation on EGFR diffusion

One key aspect not assessed directly by the studies described above is the influence of EGFR autophosphorylation on its mobility. It was previously suggested^7, 28, 31^ that EGF- induced phosphorylation of tyrosines in the EGFR C-terminal tail (C-Tail) should reduce receptor mobility by recruiting SH2 domain-containing downstream signaling molecules into a potentially large complex^1^. Groves and colleagues^59^ have further suggested that phosphorylation-dependent association of GRB2 with the EGFR C-Tail causes phase transition into a protein condensate. The EGFR slow-down seen upon EGF binding is unlikely to reflect these phenomena, since it is largely unaffected by inhibiting EGFR kinase activity with erlotinib or with the D813N mutation (in Kin^-^ EGFR). One caveat in this argument, however, is that *trans* phosphorylation of EGFR by other receptors might be responsible. We therefore investigated the effect of C-Tail phosphorylation on EGFR diffusion by collectively mutating or deleting all EGFR autophosphorylation sites.

We first mutated all 9 tyrosines in the C-tail of the receptor to phenylalanine as described by Gill *et al*.^60^, but found that this alteration severely reduced expression levels (Supplementary Figure 2). We therefore mutated these tyrosines to serines instead – retaining the polar group at each position and thus avoiding possible misfolding effects. The resulting ‘Y9S’ EGFR variant expressed at similar levels to wild type (Supplementary Figure 2a) and preserved the property of being slowed down by EGF binding (Fig. 7a and Supplementary Table 5). In the absence of EGF, the *D* value for Y9S was essentially the same as for wild type EGFR (*P* = 0.25). Adding EGF reduced *D* by 28% (*P* = 1.8 × 10^-4^) to essentially the same value (0.018 ± 0.008 μm^2^s^-1^, *P* = 1.0) as seen for EGF-bound wild type. We next also mutated a known tyrosine phosphorylation site in the TKD itself (Y845) to phenylalanine, to yield Y10X. Y845 is a SRC phosphorylation site that can also be autophosphorylated by EGFR *in vitro*^61^, and Y845 phosphorylation has been argued to induce a conformational change in the TKD^62^. The Y10X variant also showed clear evidence of EGF-induced slow-down (Fig. 7a and Supplementary Table 5), although the effect was diminished (*D* value reduced by only 19%; *P* = 0.007) as the additional Y845F mutation appeared to slow down the unliganded receptor to a significant extent (*P* = 7.9 × 10^-6^) compared with wild type and Y9S. The origin of this slow-down for Y10X is not clear, but we note that Baumdick *et al.*^62^ reported that a Y845F mutation does not affect EGFR dimerization.

**Fig. 7.**
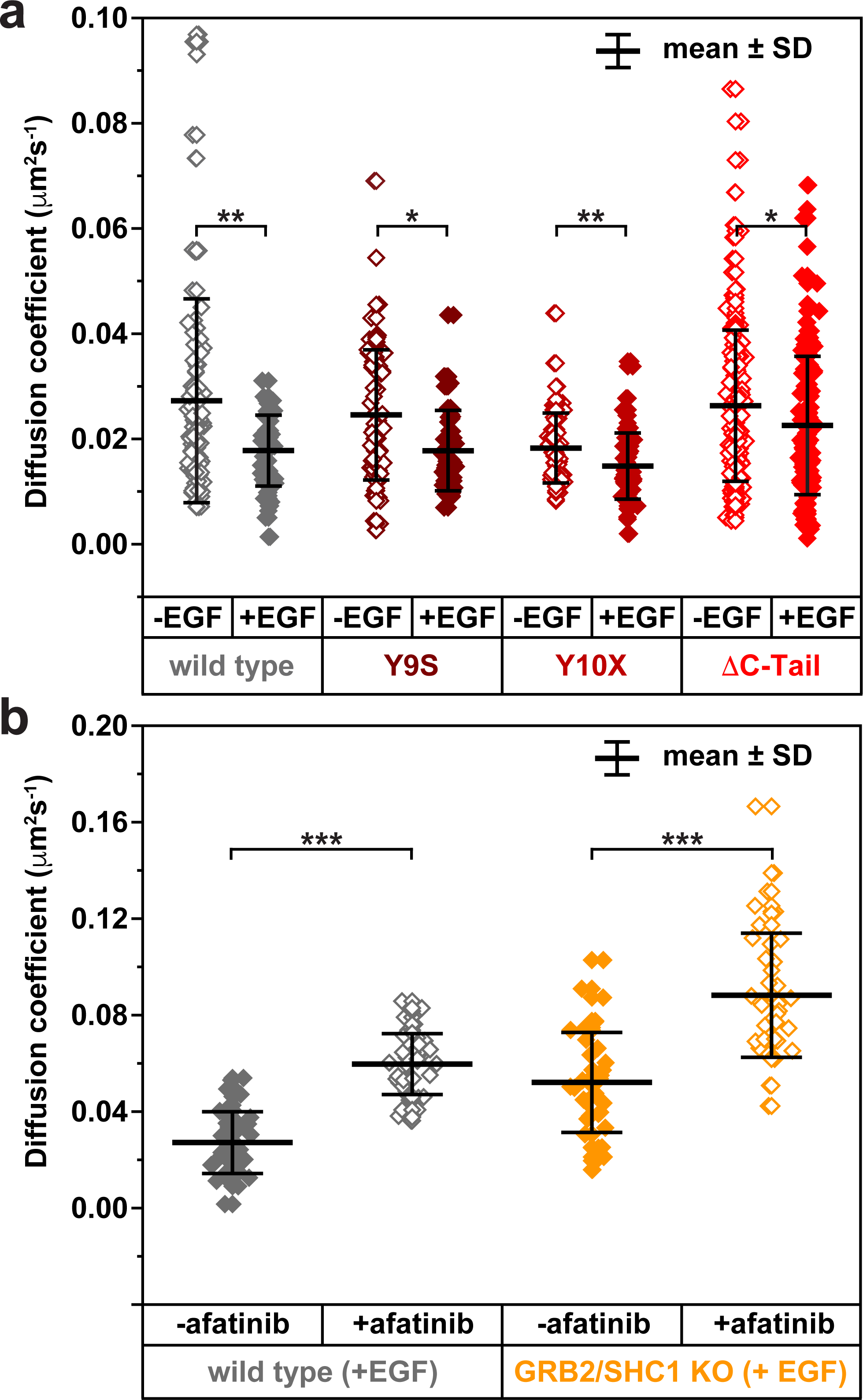
Reduced EGFR diffusion does not require C-tail phosphorylation or adaptor protein binding. **a** Mutations or truncations were made to remove phosphorylation sites from the EGFR C-tail. 200 pM QD-EGF induced a significant reduction in mean *D* value for wild type EGFR (gray) in these experiments (*P* = 4.9 × 10^-6^; *n* = 124 without ligand, 56 with ligand), as it did for variants in which either 9 (Y9s: blood red) or 10 (Y10X: medium red) phosphorylation sites were removed (Y9S: *P* = 1.8 × 10^-4^; *n* = 72 without ligand, 60 with ligand; Y10X: *P* = 0.007; *n* = 59 without ligand, 68 with ligand). Truncating the C-tail (ΔC-Tail) did not completely abolish slow-down of the receptor when activated with QD- EGF (red, *P* = 0.023; *n* = 166 without ligand, 118 with ligand), but reduced the magnitude of the effect (wild type + EGF compared with ΔC-Tail +EGF; *P* = 1.9 ×10^-3^; *n* = 56 and 118 respectively). **b** CRISPR/Cas9 was used to create a HeLa cell line in which two major adaptor proteins required for EGFR signaling – GRB2 and SHC1 – were knocked out (GRB2/SHC1 KO: see Supplementary Figure 4). Endogenous receptors were tracked using QD-EGF (200 pM) in all cases, and afatinib (10 μM) was used to reverse the effects on EGFR diffusion (see Fig. 6). QD-EGF significantly reduced EGFR diffusion in the absence of afatinib compared with in its presence in both wild type cells (gray, *P* = 6.7 × 10^-21^; *n* = 46 without afatinib, 46 with afatinib) and GRB2/SHC1 KO cells (orange, *P* = 1.6 × 10^-9^; *n* = 36 without afatinib, 42 with afatinib). Unpaired two-sided Welch’s t-tests were used to calculate *P* values (\**P* < 0.05; \*\**P* < 1×10^-3^; \*\*\**P* < 1×10^-6^), listed in Supplementary Table 5. All source data are provided as a Source data file.

We also investigated the effect of deleting the entire C-terminal tail (residues 999-1186; ΔC-Tail) on EGFR mobility (Fig. 7a and Supplementary Table 5). The C-Tail deletion did not significantly alter the *D* value in the absence of ligand (*P* = 0.65), but it did appear to blunt the EGF-induced slow-down to some extent, with EGF reducing mobility by just 14% (*D* values with EGF were 0.018 ± 0.007 μm^2^s^-1^ for wild type, and 0.023 ± 0.014 μm^2^s^-1^ for ΔC-Tail: Supplementary Table 5). This less dramatic reduction in EGF-induced slow-down is consistent with the idea that the C-Tail, especially when phosphorylated, may play a role in promoting intracellular inter-receptor interactions in the ligand-occupied receptor. By using FRET-FLIM to assess dimerization of this construct, we further confirmed that the reduced EGF-induced slow-down of ΔC-Tail EGFR does not reflect an inability to dimerize (Supplementary Figure 3).

Finally, in a complementary approach to address the potential role of adaptor protein recruitment in the EGF-induced slow-down of EGFR, we created HeLa cell lines in which CRISPR was used to knock out two key adaptor proteins in EGFR signaling: SHC1 and GRB2 (Supplementary Figure. 4). As shown in Fig. 7b and Supplementary Table 5, loss of SHC1 and GRB2 did not prevent the EGF-induced slow-down of EGFR. Since we studied endogenous EGFR in these cells, we could not measure unliganded receptor diffusion directly. Instead, we assessed the ability of afatinib to increase mobility of QD-EGF-bound EGFRs. Just as afatinib prevents EGF from slowing down EGFR (Fig. 6), so does it speed up EGF-bound EGFR in parental HeLa cells (Fig. 7b, left). Afatinib also significantly increased *D* for EGF-bound EGFR in GRB2/SHC1 knock- out HeLa cells, as shown in Fig. 7b, right (*P* = 1.6 × 10^-9^; Supplementary Table 5). This finding argues against a role for signaling molecule recruitment or protein condensation mediated by multivalent signaling adaptor proteins in reducing EGFR diffusion upon EGF stimulation.

### EGFR slow-down correlates with EGFR clustering

The SPT data presented here argue that, although necessary, receptor dimerization alone is not sufficient for the observed slow-down when EGF binds intact wild type EGFR in cells. Whereas EGF-induced dimerization appears to be driven primarily by the ECR, slow-down of EGFR diffusion requires additional key interactions related to asymmetric TKD dimerization (involving both TKD and JM regions), with additional possible contributions from the C-Tail. Several studies have noted that the EGFR TKD forms a ‘daisy chain’ of N-lobe-to-C-lobe asymmetric dimers in crystals^6, 9, 12^. These interactions lead to precipitation of the protein *in vitro* when the JM region is included^9^. One intriguing possibility is that ligand-induced receptor dimers associate with one another into larger oligomers by forming such TKD daisy chains, with additional contributions from the ECR as has been suggested^24^. Another model for dimers of dimers also requires that the TKD is in the active conformation^25^, but differs substantially in detail. To test the hypothesis that the ligand-induced EGFR slow-down reflects formation of higher-order oligomers, we used direct stochastic optical reconstruction microscopy (dSTORM) super-resolution imaging of HA-tagged EGFR with a AlexaFluor647-labeled HA tag antibody. As shown in Fig. 8a, both wild type EGFR and ΔICR EGFR are quite evenly distributed across CHO cells in the absence of EGF. When EGF is added, however (at 50 nM), significant clustering of the wild type receptor, but not ΔICR EGFR is seen (Fig. 8a). The difference in clustering can be seen by comparing a cumulative distribution function (CDF) plot of cluster sizes (Fig. 8b).

**Fig. 8.**
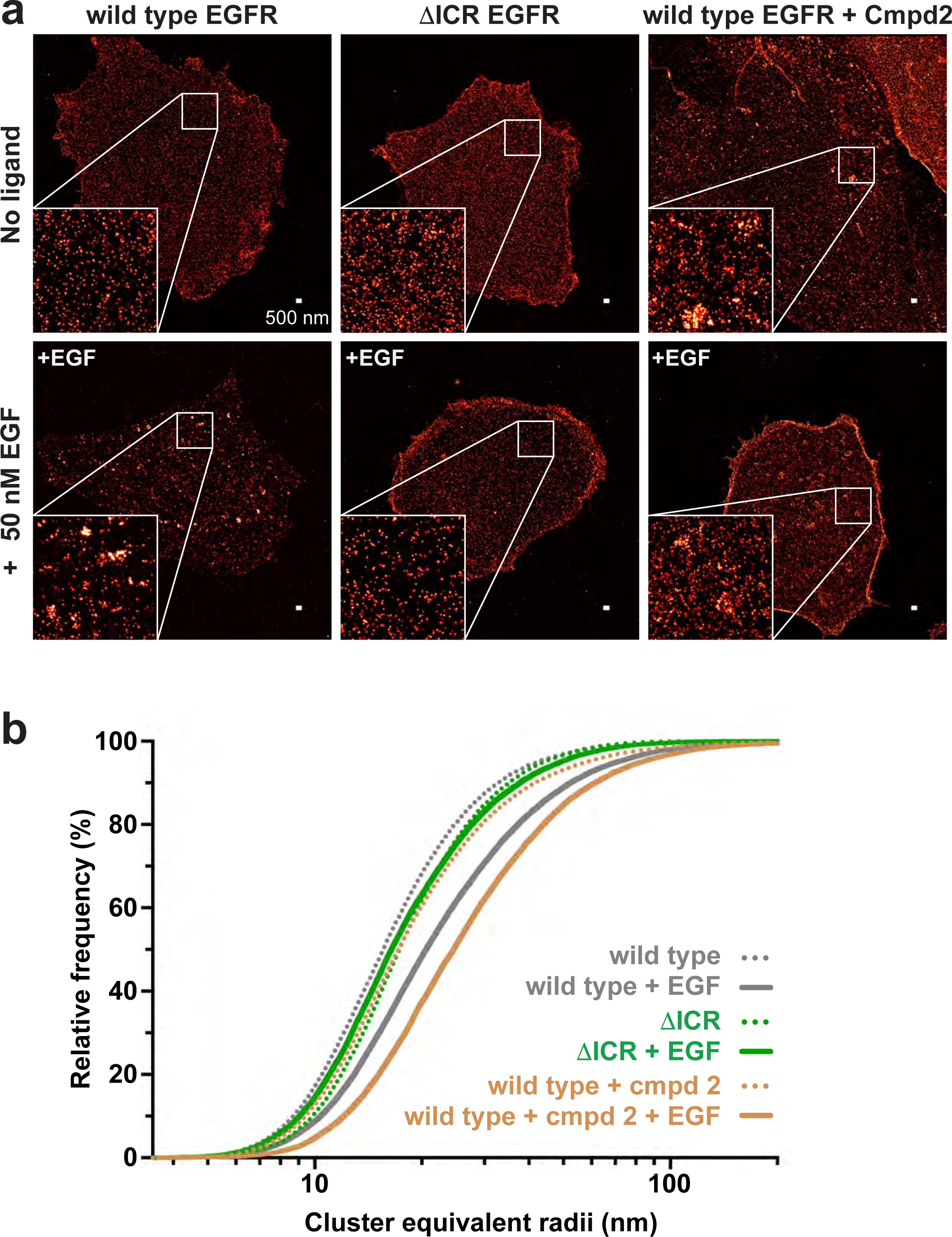
Clustering of EGFR on the cell membrane correlates with the slow-down of receptor diffusion. **a** Using dSTORM super-resolution imaging, three distinct conditions were visualized and compared. CHO cells expressing wild type EGFR exhibited clustering upon ligand stimulation (50 nM EGF). In contrast, the ΔICR construct, which did not slow down with EGF activation in SPT experiments, showed no evidence of clustering when stimulated with saturating levels of EGF. CHO cells expressing wild type EGFR and treated with compound 2 (10 μM) showed a limited degree of clustering in the absence of EGF that was significantly increased upon EGF addition. **b** DBSCAN was used to quantify EGFR cluster size from dSTORM images, as described in Methods. Cluster sizes are represented as the cumulative distribution of cluster equivalent radii per condition (in nm), and a shift to the right indicates larger clusters. Treatment with EGF induces the formation of larger EGFR clusters (solid gray curve) when compared with untreated cells (dotted gray curve). The results also show that compound 2 induces some clustering (dotted gold curve) that is substantially further increased upon EGF treatment (solid gold curve) – with EGF and compound 2 together promoting the greatest degree of clustering seen in the study. Untreaded ΔICR-EGFR shows a slightly larger degree of clustering than expected (dotted green curve), but is unaffected by EGF addition (solid green curve). A Kruskal-Wallis one way ANOVA test with post-hoc correction for multiple comparisons (Dunn’s test) was used to confirm that all conditions differ significantly from untreated wild type EGFR (*P* < 0.0001).

This result supports the suggestion from the SPT data that the ECR is sufficient for ligand-induced EGFR dimerization (which is retained for ΔICR EGFR), but that the ICR is required for formation of higher order oligomers. Importantly, adding the compound 2 kinase inhibitor further increased clustering of EGF-bound EGFR (Fig. 8a,b), presumably by further stabilizing the active TKD conformation. Compound 2 alone also shifted the CDF plot for wild type to the right even in the absence of EGF (*P* < 0.0001), consistent with its ability to slow down the receptor without extracellular ligand as shown in Fig. 6. These results suggest that wild type EGFR is driven into micron-sized clusters only when the TKD is caused to adopt the active conformation. Ordinarily, this occurs upon EGF binding to the ECR, but it can be mimicked – without any tyrosine phosphorylation – when compound 2 helps drive the wild type TKD into an active dimeric conformation.

## Discussion

Single-particle tracking has been used to study the dynamics of EGFR for over two decades^2, 7, 27, 29, 30, 63, 64^, with the commonly observed reduction in diffusion coefficient frequently being equated with receptor dimerization and signaling. We confirmed that the EGFR slow-down requires ligand-induced dimerization, and set out to ask which domains within the receptor dominate in driving this process. We found that changes in EGFR diffusion correlated with altered ECR dimerization strength^18, 30^, but were unaffected by the loss of putative dimerization motifs in the transmembrane region, consistent with previous studies^42, 43^. Importantly, removing the entire ICR left ligand-induced EGFR dimerization intact but abolished the ligand-induced receptor slow-down. Thus, our results suggest that EGFR dimerization and the ligand-induced receptor slow- down seen in SPT studies are distinct phenomena. Across a series of EGFR variants that retain the (ECR-driven) ability to dimerize upon EGF binding, we further showed that the ligand-induced EGFR slow-down does not depend on kinase activity. However, it requires both intracellular JM interactions and N-lobe/C-lobe TKD interactions associated with the asymmetric dimer formed by EGFR TKDs in their active conformation. The C-Tail also appears to make a positive contribution.

What is responsible for the reduced diffusion of EGF-bound EGFR? Activated RTKs are well known to recruit multivalent signaling adaptor molecules after tyrosine autophosphorylation^1^. These molecules can undergo protein condensation or phase transitions^65^ that have recently been argued to be important for RTK signaling^59, 66^. It does not appear that this phenomenon can explain ligand-induced EGFR slow-down, since it was not affected by a loss of kinase activity, by mutating all EGFR autophosphorylation sites, or by loss of the signaling adaptor molecules SHC1 and GRB2. One possibility is that the reduced diffusion seen in our SPT studies reflects formation of discrete higher-order EGFR oligomers, as have been suggested and modeled by others^21, 26^. Needham *et al.*^25^ argued that the TKD is key for forming tetramers or higher order oligomers of EGFR that they deduced using a method called fluorophore localization imaging with photobleaching (FLImP). This group proposed a model in which EGFR dimers associate with one another both through a set of ECR interactions and by dimerization of the asymmetric TKD dimer. Huang *et al.*^24^ also combined single molecule imaging and MD studies to support a model for EGF-bound EGFR tetramers and larger oligomers that is related conceptually, but is very different in structural detail. They proposed that a region of domain IV drives association of ECR dimers, and that the TKD forms daisy chains of the sort seen in crystals. Although they have yet to be defined in detail, oligomers of this general class could be responsible for both the EGF-induced slow-down of EGFR in SPT studies and the EGFR clusters seen by dSTORM. It is also possible the other molecules or scaffolds are involved in the observed clusters. In either case, we suggest that the structurally well-characterized ECR dimerization interface^14^ is the major interaction site that ‘switches’ EGFR between states, given that the ECR dimerizes strongly enough when bound to EGF (*K*_d_ ∼1 μM) to drive the majority of receptors in a cell into dimers. The other interactions involved in clustering are much weaker, including those between EGFR intracellular regions – for which we estimate *K*_d_ >100 μM from our studies of the JM-TKD region (Fig. 4d) – and other proposed ECR interactions that remain to be characterized. It seems highly likely that higher-order receptor assembly/clustering involves cooperation of multiple weak interactions involving the ECR, the ICR, and possibly the C-Tail (consistent with other suggestions^67^).

Our observations are also related to findings that simply dimerizing EGFR with a bivalent antibody^68^ or chemical biology tool^69^ is sufficient to promote receptor autophosphorylation and GRB2 recruitment, but fails to activate ERK signaling. Artificially dimerized EGFR also fails to form the clusters seen when EGFR is activated with EGF itself^69^. In a related vein, the Jura lab concluded from STORM imaging studies that EGFR clustering depends on its ability to form a catalytically active complex^70^, a finding that is underlined by our studies with EGFR inhibitors that stabilize different TKD ‘states’. Taken together, the observations made here provide a platform for dissecting mechanisms of EGFR oligomerization that we expect will provide important insights into how this complex receptor signals, and may illuminate new avenues for therapeutic interventions when it signals inappropriately.

## Methods

### Reagents

Biotinylated anti-HA fab fragments were purchased from Roche (#12158167001), and EGF biotinylated at a 1:1 stoichiometry was purchased from Life Technologies (# E- 3477). Streptavidin-conjugated quantum dots (QD) were purchased from Life Technologies (QD-605 #Q10101MP or QD-655 #Q10121MP). JaneliaFluor dyes JF-549 and JF-646 were a gift from the Luke Lavis Lab. Afatinib and erlotinib were obtained from SelleckChem (#S1011 and #S7786 respectively), and compound 2 was synthesized in house as described^57^. All inhibitors were dissolved at 10 mM in DMSO for stock solutions and diluted directly into culture medium for imaging (alongside DMSO-only control).

### Tissue culture and transfection

CHO cells (ATCC) were grown in complete DMEM/F12 medium (ThermoFisher, #11320033) containing 10% fetal bovine serum (ThermoFisher) and 1% penicillin-streptomycin (ThermoFisher, #15140148) at 37°C, in a humidified incubator with 5% CO_2_. Cell transfection via electroporation used a Nucleofector 2b (Lonza), following the manufacturer’s protocol. Immediately after electroporation, ∼0.5 ml of prewarmed antibiotic-free complete media was added to the electroporation cuvette and the cell suspension was transferred to 35 mm poly-D-lysine coated FluoroDish optical dishes (WPI, #FD35PDL-100) for a final volume of 2 ml of antibiotic-free complete media for test imaging, or to 25 cm flasks to generate stable cell lines. Optical dishes were cleaned using 1M KOH for 30 min, then rinsed with Milli-Q H_2_O for 5 min, then incubated with 100% ethanol for 30 min, followed by another 5 min rinse with Milli-Q H_2_O. For generation of stable cell lines, cells were selected in complete media containing 0.5 mg/ml G418 (Gibco, #10131035). HeLa were cultured in DMEM (Life Technologies, Carlsbad, CA; #10313-021) supplemented with 10% fetal bovine serum (HyClone, Logan, UT), penicillin–streptomycin, and 2 mM L-glutamine.

### Plasmids and cloning

Human EGFR was tagged with a hemagglutinin tag to yield HA-EGFR as previously described^31^, and all subsequent EGFR constructs were made using the HA-EGFR plasmid and Platinum SuperFi II PCR Master Mix (Invitrogen, #12368010). Halo-EGFR constructs used for FRET-FLIM experiments were created using the HA-EGFR plasmid and the Gibson assembly approach to replace the N-terminal HA tag with the HaloTag® plasmid (Promega, #G7711). The final resulting construct contained the HaloTag and a GSGS linker region after the EGFR signal sequence. ‘Y9F’, ‘Y9S’, and ‘Y10X’ EGFR constructs were made using Gibson assembly of nine overlapping dsDNA fragments. All DNA fragments were generated by PCR using Q5 Hot Start High-Fidelity DNA polymerase (NEB # M0494S). The primers for each fragment had gene-specific 3’- sequences and 15–20 additional bases at the 5’ end that overlap with the ends of the adjacent fragment. PCR products were purified by QIAquick Gel Extraction Kit (Qiagen # 28704) and used in equimolar amounts in a Gibson assembly reaction with NEBuilder HiFi DNA Assembly Master mix (NEB # E2621S). The reaction was incubated at 50°C for 60 min and directly transformed into chemically competent NEB 10-beta *E. coli* cells (NEB #C3019H) and plated onto LB agar plates. Multiple colonies were screened by Sanger sequencing to identify mutated constructs.

The TM3X construct was made in two steps. In the first, T624V, G625V, G628V, and A629V mutations were introduced simultaneously with a set of primers. The resulting construct was then used as a template to introduce A637V and G641V mutations. The JM4X construct was generated in a similar sequential manner. The ΔICR and ΔC-tail constructs were generated using primers with ∼15 nucleotides overlapping the desired ending region (ending at 670 and 998 respectively) followed by an introduced stop-codon. All constructs were sequence verified prior to transfection.

### Analysis of EGFR signaling and signaling molecule expression by immunoblotting

For cell signaling studies, CHO cells expressing the relevant EGFR variants were serum-starved overnight and either left unstimulated or stimulated with 16 nM EGF (R & D Systems) for 5 min. Total cell lysates were prepared and analyzed by SDS-PAGE using 4-12% gradient Bis-Tris NuPAGE gels (ThermoFisher) followed by immunoblotting^71^. For signaling studies, primary antibodies were all used at a 1:1,000 dilution as follows: for phosphorylated EGFR (pY1173: CST#4407), total EGFR (R&D AF231) and ERK1/2 (pT202/pY204: CST #9106; total ERK: CST#4696). Secondary antibodies were horse anti-mouse IgG horseradish peroxidase (HRP)-linked antibody (CST #7076) used at 1:10,000 dilution, WestVision anti-rabbit IgG (H+L) HRP polymer (Vector Labs WB-1000) used at 1:10,000, and rabbit anti-goat IgG conjugated to HRP (R&D HAF017) used at 1:1,000 dilution. Chemi-luminescence signals were detected using SuperSignal West Pico Chemiluminescent Substrate (ThermoFisher Scientific), visualized and quantified using a Kodak Image Station 440CF (Kodak Scientific). Blotting for GRB2 (1:1,000; CST #3972) was used as a loading control^72^.

For analysis of GRB2, SHC1, and EGFR expression in cells after CRISPR knock-out, antibodies were again used at 1 1:1,000 dilution. Rabbit Polyclonal (C-23: SantaCruz #SC-255) was used for GRB2, mouse monoclonal (BD Biosciences #610878) for SHC1, a β-actin mouse monoclonal (Sigma Aldrich #A1978) as loading control, and a rabbit monoclonal antibody against EGFR (Cell Signaling Technology EGF Receptor (D38B1) mAb #4267, used at 1:2,000 dilution). Detection was as above or employed (for LI-COR) LI-COR BioScience IRDye 800CW Goat Anti-Rabbit IgG #926-32211 as secondary antibody at a 1:20,000 dilution in 3% BSA/PBS for 1 h at room temperature.

### Single-particle tracking (SPT)

Single-particle was carried out using a Nikon Ti-2 N-STORM/TIRF inverted microscope equipped with a 100x 1.49 NA oil immersion objective with objective heater and Tokai Hit stage-top incubator, both kept at ∼37°C. Excitation was provided by a 405 nm laser at 3.2 mW power. Emission was detected using an Andor iXon 897 EMCCD (pixel size 158.15 nm at 46 Hz) using a Hamamatsu Gemini W-View dual image splitter for one- camera 2-color imaging, with a 600/40 nm BP filter, 655 nm LP filter, and a T640LPXR- UF2 dichroic beam splitter from Chroma. For SPT of QD-tagged HA-EGFR^31^, biotinylated anti-HA or EGF were conjugated to streptavidin-tagged quantum dots by incubating equimolar mixtures of either anti-HA Fabs or EGF with the QDs in phosphate-buffered saline (PBS) plus 1% bovine serum albumin (BSA) at 4°C for 2 h with agitation before imaging. CHO cells stably expressing HA-EGFR were plated the day before imaging in 35 mm FluoroDishes using complete medium, and then incubated with complete DMEM/F12 medium that lacked FBS (starvation medium) for 6 to 12 h prior to imaging experiments. For experiments using TKIs, cells were incubated with the appropriate TKI at a concentration of 10 μM for 30 min at 37°C in starvation medium. For all imaging experiments, cells were then incubated with 200 pM anti-HA-QD605/655 in 1 ml of starvation medium for 5 min at 37°C or with 200 pM EGF-QD605/655 in 1 ml of starvation medium for 1 min at 37°C. Excess dye was extensively washed out using PBS. Cells were then imaged in ‘Live Cell Imaging Solution’ (Invitrogen, #A14291DJ) supplemented with 10 mM glucose at 37°C. Each cell was imaged for 1 min, and each optical dish was subjected to imaging for no more than 10 min.

### Receptor tracking

Localizations were generated from raw data movies using a 2D localization algorithm based on maximum likelihood estimation assuming a 2D Gaussian point-spread function (PSF) model^73^. Thresholds on photons, localization error, PSF width, and goodness-of-fit as defined by a *P* value were used to remove low-quality and false-positive localizations.

SPT was performed using custom-written software in MATLab, implementing a modified version of the algorithm presented by Jaqaman *et al*.^74^. The algorithm was modified by defining physically accurate cost functions for each of the state assignments, accounting for fluorophore photophysics and emitter diffusion. Briefly, we define the costs by assuming a two-state fluorophore kinetic model (on or off), Brownian diffusion of the emitters, and Gaussian localization errors. With the redefined costs in both frame-to-frame and gap-closing cost matrices, our algorithm otherwise proceeds similar to that described by Jaqaman and colleagues^74^. Single component diffusion coefficients were estimated from SPT results as follows: The mean squared displacement (MSD) was computed for non-overlapping time lag for all individual trajectories, after which the ensemble MSD was calculated for each cell. The first 5 points of the MSD versus time lag were fit to a linear (Brownian) motion model using weighted least-squares, where the weights were chosen to be the inverse of the variance of each point of the MSD data (i.e., the weights are given as *N*/Δ*t*^2^, where *N* is the number of displacements used to compute the MSD at a time lag of Δ*t*). The diffusion coefficient is then taken to be *m*/4, where *m* is the slope of the MSD. Data outputs from MATlab were graphed and analyzed using Origin 2021b (9.8.5.201).

### FRET-FLIM imaging

FRET-FLIM experiments were carried out using a Leica Stellaris 8 Falcon system with a 100x 1.4 NA oil immersion objective. CHO cells were stably transfected with Halo-EGFR constructs and seeded into FluoroDishes in complete medium one day prior to imaging. Cells were then grown in complete medium lacking FBS for 6-12 h before labeling with dyes. CHO cells were then incubated with 20 nM JF-549, JF-646 or an equimolar concentration of both dyes in 1 ml of complete media for 5 min at 37°C. Excess dye was extensively washed out using PBS, and cells were imaged in Live Cell Imaging Solution (Invitrogen, #A14291DJ) supplemented with 10 mM glucose at 37°C. Each cell was imaged until 100 photons were collected per pixel for FRET-FLIM analysis. Using the analysis software package on the Stellaris 8 Falcon, lifetimes (τ) were calculated using the following formula:

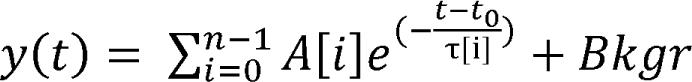

Where *n* is the number of exponential components, *t_0_*is the lifetime offset, *A* is the amplitude, τ is the exponential decay (lifetime), and *Bkgr* is the background correction. Data outputs from the Stellaris 8 Falcon software package were graphed and analyzed using Origin 2021b (9.8.5.201).

### Protein Purification for AUC studies

DNA encoding JM-TKD [645-998] and TKD [672-998] protein regions was subcloned into pFastBac (ThermoFisher Scientific) following a hexa-His N-terminal tag. Agilent site-directed mutagenesis was used to introduce the V924R mutation into JM-TKD and the I682Q mutation into TKD (using numbering for mature protein). Protein expression employed the Bac-to-Bac system (Invitrogen) exactly as previously described^75^, using a Ni-NTA column followed by anion exchange chromatography and gel filtration.

Extinction coefficients for both JM-V924R-TKD and I682Q-TKD were determined to be 52,370 M^-1^ cm^-1^ using the ExPASy ProtParam Tool^76^. Purification of the histidine-tagged EGFR extracellular region was performed exactly as described in previous studies^51^.

### Sedimentation Equilibrium Analytical Ultracentrifugation (SE-AUC)

Freshly purified JM-V924R-TKD and I682Q-TKD proteins were mixed in a 1:1 molar ratio at three total protein concentrations (28 μM, 18 μM, and 10 μM) and centrifuged at three speeds (9,000 rpm, 12,000 rpm, and 15,000 rpm) at 4°C until equilibrium was reached using an An Ti 60 rotor in an XL-I analytical ultracentrifuge (Beckman), run using ProteomeLab XL-A/XL-I (version 6.2). SE-AUC experiments using the EGFR extracellular region (sEGFR) used sEGFR at concentrations of 10 μM, 5 μM, and 2 μM, with- and without addition of TGFα at a 1.2-fold molar excess, as previously described^46^. In all experiments, radial measurements of absorbance at 280 nm (A_280_) and 289 nm (A_289_) were taken using step scan(s) from 5.8 – 7.3 cm at 0.003 cm intervals, with 5 replicates. Data points in Fig. 4c,d are raw AUC centrifugation data plotted as the natural logarithm (ln) A_280_ against a function of radius squared (*r*^2^ - *r* ^2^)/2, where *r* is the radial position in the sample and *r*_o_ is the radial position of the meniscus. For a single ideal species, this representation gives a straight line with slope proportional to its molecular mass. Straight lines plotted in Fig. 4c,d correspond to expected results for monomer (gray) or dimer (red) of sEGFR (Fig. 4c) or for monomer (gray) or dimer (red) of the kinase domain (Fig. 4d). Estimates of dimerization dissociation constant (*K*_d_) for sEGFR were obtained from more complete analysis, fitting data for three concentrations and three speeds to a model describing simple dimerization of a 1:1 sEGFR:TGFα complex:

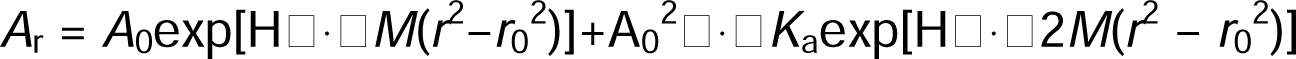

where *A*_r_ is the absorbance at radius *r*, *A*_0_ is the absorbance at the reference radius *r*_0_, *M* is the molecular weight of the 1:1 complex), H is the constant [(1 – ∇ρ)ω^2^]/2RT, ∇ is the partial specific volume (estimated at 0.73 ml/g), ρ is the solvent density (1.003 g/ml), ω is the angular velocity of the rotor (radians/sec), R is the gas constant, T is absolute temperature, and *K*_a_ is the fitted parameter corresponding to the equilibrium constant for dimerization (1/*K*_d_) as described^46^. Experiments using JM-V924R-TKD and I682Q-TKD, showed no evidence for dimerization, so could not be fit. Data were plotted and analyzed using GraphPad Prism 9 for MacOS (Version 9.5.1).

### Genome editing in HeLa cells

Cas9-mediated DNA cleavage was used to knock out the endogenous genes encoding GRB2 and SHC in HeLa-CCL2 cells by splicing out the exon containing the SH2 domain of each respective gene. Three guide RNAs (gRNAs) targeting either the fourth exon of human SHC1 (5’-CCCGCTCAGCTCTATCCTGG-3’), (5’- CGCCCGCTCAGCTCTATCCT-3’), (5’-TGTAGCCGCCCGCTCAGCTC - 3’) or the third exon of human GRB2 (5’-CTTAGCAAACAGCGGCACGA-3’), (5’- CAGAGCCAAGGCAGAAGAAA-3’), (5’-TGGCAAAATCCCCAGAGCCA-3’) were designed using the http://crispr.mit.edu portal. The gRNAs were subcloned into *Bbs I*- digested PX458 vector (Addgene plasmid #48138) using Gibson assembly. This generated a plasmid that could be used for simultaneous expression of the gRNA, WT Cas9, and a green fluorescent protein (GFP) reporter. The plasmid was used to transiently transfect HeLa-CCL2 cells using the Amaxa system (Lonza) with Solution R and Program I-013. Flow cytometry (iCyt cell sorter) was used to select GFP-positive cells, which were immediately plated at single-cell density in 96-well plates. Successful knock-out of the targeted proteins was then screened for using immunoblotting (see Supplementary Figure. 4) and was confirmed with at least two independent sample preparations.

### Super-resolution dSTORM imaging

CHO-K1 cells stably expressing HA-tagged wild type or ΔICR EGFR were plated overnight on piranha-cleaned 25 mm coverslips. Cells were serum starved for at least 2 h at 37°C. EGF-stimulated cells were treated with 50 nM recombinant human EGF (PeproTech, #AF-100) for 3 min at 37°C to ensure activation of EGFR on the basal surface of the cell. Where noted, cells were pre-treated with 10 μM of the Compound 2 kinase inhibitor for 30 min in serum-free medium. Cells were washed with PBS and then fixed with 100% MeOH at 20°C for at least 30 min, rehydrated with PBS, blocked with 3% BSA/PBS, and labeled with AlexaFluor647-labeled anti-HA.11 Epitope Tag antibody (BioLegend, #16B12) at 5 μg/ml for 2 h at room temperature. Cells were then washed and post-fixed with 4% paraformaldehyde for 15 min at room temperature and washed twice with PBS and once with 10 mM Tris-HCl (pH 7.2) and stored in PBS until imaging. Imaging was performed in a standard dSTORM imaging buffer with an enzymatic oxygen scavenging system and primary thiol: 50 mM Tris, pH 8.5 containing 10 mM NaCl, 10% w/v glucose, 168.8 U/ml glucose oxidase (Sigma #G2133), 1404 U/ml catalase (Sigma #C9322), and 60 mM 2-aminoethanethiol (MEA). Data were collected using a custom-built microscope^77^ consisting of a 647 nm laser excitation source (500 mW 2RU-VFL-P; MPB Communications Inc.), an sCMOS camera (C11440-22CU; Hamamatsu Photonics) a 1.35 NA silicon oil immersion objective (Olympus UPLSAPO100XS) and a 708/75 nm emission filter (FF01-708/75-25; Semrock). A total of 60,000 frames were collected per cell with an exposure time of 50 ms per frame. Images were registered every 10,000 frames^77^.

### Super-resolution image reconstruction and data analysis

dSTORM data were analyzed using custom-written software (SMITE (https://github.com/LidkeLab/smite). Localizations with intensities greater than 4000 photons were removed from the lists of coordinates. Analysis of dSTORM EGFR clustering was performed using the density-based DBSCAN algorithm^78^ implemented in MATLab^79, 80^. Parameters chosen were a maximal distance between neighboring cluster points of ε = 20 nm and a minimum cluster size of 10 observations. Cells were segmented into 2 μm × 2 μm regions of interest (ROIs) for analysis. To avoid potential influence of receptor density/expression levels on clustering results, we removed from analysis all ROIs with >25,000 localizations per ROI in order to maintain similar densities between conditions. Cluster boundaries were produced with the MATLab ‘boundary’ function using a default methodology that produces contours halfway between a convex hull and a maximally compact surface enclosing the points. The cluster areas within these boundaries were then converted into the radii of circles of equivalent area for a more intuitive interpretation. A Kruskal-Wallis one way ANOVA test was conducted to examine the differences between the sizes of cluster equivalent radii calculated from DBSCAN cluster analysis. Post-hoc correction for multiple comparisons (Dunn’s test) between the cluster sizes of wild type untreated EGFR and other conditions were additionally conducted as shown in Fig. 8b.

#### Data availability

Data supporting the findings of this study are all available within the article and Supplementary Information files. All cell lines and plasmids generated for the work in this manuscript are available upon request from the authors. PDB entry 3GOP was used in generation of Fig. 5a. Source data are provided with this paper, including all uncropped gels. Custom-written software for analyzing dSTROM data (SMITE) is available from the following link: https://github.com/LidkeLab/smite. Requests for materials and other correspondence should be addressed to Mark A. Lemmon, Krishna C. Mudumbi, or Diane S. Lidke.

## Acknowledgements

We thank members of the Lemmon and Lidke laboratories and Prof. Kathryn Ferguson for comments on the data and the manuscript. This material is based on work supported in part by the National Institutes of Health (R01-CA248166 to M.A.L. and D.S.L., K99- CA256250 to K.C.M.), an Arnold O. Beckman Postdoctoral Fellows Award (17-001959, to K.D.A.). We thank Dr. Cédric Cleyrat for generating the SHC/GRB2 knock-out cells, Drs. Mara Steinkamp and Carolina Nitto Franco for generating initial EGFR mutations and cell lines, and Dr. Michael Wester for assistance with dSTORM imaging analysis. We gratefully acknowledge use of the Yale West Campus Imaging Core and the University of New Mexico Comprehensive Cancer Center fluorescence microscopy and flow cytometry facilities, which are supported at UNM by NIH grant P30-CA118100.

## Author contributions

K.C.M., D.S.L. and M.A.L. conceived the overall project. K.C.M. was responsible for FRET-FLIM studies and most SPT analyses described here, with assistance from L.W.K. and E.M.M.. Other SPT analyses were performed by I. O.-C. and E.A.B. Z.O.P. was responsible for biophysical studies of the EGFR ECR and TKD. A.K. performed cell signaling studies, with assistance from K.C.M., L.W.K., C.H., and H.R.O.. E.A.B performed dSTORM imaging and analysis. K.D.A. synthesized compound 2 for this study. Imaging analysis methods were developed by D.J.S. and K.A.L.. K.C.M., D.S.L., and M.A.L. supervised the overall project and wrote the manuscript, on which all authors commented.

