## Supplementary Information for "Distinct interactions stabilize EGFR dimers and higher-order oligomers in cell membranes"

**Supplementary Tables 1-5 and Supplementary Figures 1-4**

**Supplementary Table 1.**

SPT diffusion results for intact EGFR with N-lobe (I682Q) and C-lobe (V924R) TKD mutations. Measurements in the absence of ligand used QD-conjugated HA tag antibody Fab fragments to label the receptor. Measurements with EGF employed QD-EGF at 200 pM.

| Condition 1 | *D* ($\mu^2 s^{-1}$) | Condition 2 | *D* ($\mu^2 s^{-1}$) | % Change | *P* value | $n_1$ | $n_2$ |
|---|---|---|---|---|---|---|---|
| WT no EGF | 0.025±0.009 | WT + EGF | 0.018±0.008 | 27.6% | $1.5\times10^{-9}$ | 126 | 118 |
| I682Q no EGF | 0.025±0.013 | I682Q + EGF | 0.028±0.016 | 11.9% | 0.13 | 116 | 116 |
| V924R no EGF | 0.023±0.010 | V924R + EGF | 0.022±0.011 | 5.48% | 0.36 | 108 | 124 |

**Supplementary Table 2.**

SPT diffusion results for EGFR harboring extracellular glioblastoma mutations, unstimulated of saturated with EGF or EREG. QD-conjugated HA tag antibody Fab fragments were used to label the receptor both in the absence and presence of ligand. Unlabeled EGF was added at 50 nM, and unlabeled EREG was added at 1 μM.

| Condition 1 | *D* ($\mu^2 s^{-1}$) | Condition 2 | *D* ($\mu^2 s^{-1}$) | % Change | *P* value | $n_1$ | $n_2$ |
|---|---|---|---|---|---|---|---|
| WT no ligand | 0.022±0.012 | WT + EGF | 0.012±0.008 | 43.6% | $5.5\times10^{-6}$ | 67 | 33 |
| WT no ligand | 0.022±0.012 | WT + EREG | 0.017±0.010 | 22% | 0.044 | 67 | 26 |
| R84K no ligand | 0.019±0.009 | R84K + EGF | 0.0095±0.007 | 50.2% | $1.7\times10^{-6}$ | 35 | 38 |
| R84K no ligand | 0.019±0.009 | R84K + EREG | 0.011±0.006 | 42.7% | $6.0\times10^{-5}$ | 35 | 27 |
| A265V no ligand | 0.018±0.006 | A265V + EGF | 0.013±0.005 | 29.0% | $3.8\times10^{-4}$ | 36 | 31 |
| A265V no ligand | 0.018±0.006 | A265V + EREG | 0.012±0.004 | 30.8% | $1.4\times10^{-4}$ | 36 | 21 |
| WT + EGF | 0.012±0.008 | WT + EREG | 0.017±0.010 | 27.7% | 0.047 | 67 | 26 |
| R84K + EGF | 0.0095±0.007 | R84K + EREG | 0.011±0.006 | 14.9% | 0.38 | 38 | 27 |
| A265V + EGF | 0.013±0.005 | A265V + EREG | 0.012±0.004 | 2.53% | 0.80 | 31 | 21 |

**Supplementary Table 3.**

SPT diffusion results for EGFR with mutations in the transmembrane domain (TM3X), juxtamembrane domain (JM4X), and a construct where the kinase domain has been removed (ΔICR). Measurements in the absence of ligand used QD-conjugated HA tag antibody Fab fragments to label the receptor. Measurements with EGF employed QD-EGF at 200 pM.

| Condition 1 | $D$ (μ$^2$s$^{-1}$) | Condition 2 | $D$ (μ$^2$s$^{-1}$) | % Change | $P$ value | $n_1$ | $n_2$ |
|---|---|---|---|---|---|---|---|
| WT no EGF | 0.019±0.007 | WT + EGF | 0.012±0.006 | 37.1% | $6.7\times10^{-6}$ | 40 | 42 |
| TM3X no EGF | 0.019±0.010 | TM3X + EGF | 0.009±0.006 | 52.2% | $1.9\times10^{-7}$ | 45 | 46 |
| WT no EGF | 0.016±0.008 | WT + EGF | 0.010±0.005 | 37.5% | $9.3\times10^{-4}$ | 32 | 34 |
| JM4X no EGF | 0.015±0.006 | JM4X + EGF | 0.015±0.007 | 1.39% | 0.91 | 24 | 30 |
| WT no EGF | 0.028±0.023 | WT + EGF | 0.018±0.007 | 36.4% | $9.2\times10^{-6}$ | 124 | 56 |
| ΔICR no EGF | 0.026±0.013 | ΔICR + EGF | 0.028±0.013 | 7.87% | 0.21 | 116 | 121 |

**Supplementary Table 4.**

SPT diffusion results for kinase-dead EGFR and wild type EGFR treated with different inhibitors. Measurements in the absence of ligand used QD-conjugated HA tag antibody Fab fragments to label the receptor. Measurements with EGF employed QD-EGF at 200 pM.

| Condition 1 | $D$ (μ$^2$s$^{-1}$) | Condition 2 | $D$ (μ$^2$s$^{-1}$) | % Change | $P$ value | $n_1$ | $n_2$ |
|---|---|---|---|---|---|---|---|
| No drug no EGF | 0.020±0.008 | No Drug + EGF | 0.012±0.006 | 38.5% | $1.0\times10^{-6}$ | 39 | 63 |
| Kin$^-$ no EGF | 0.021±0.006 | Kin$^-$ + EGF | 0.016±0.006 | 24.2% | $1.2\times10^{-4}$ | 48 | 46 |
| Erlotinib no EGF | 0.018±0.009 | Erlotinib + EGF | 0.011±0.007 | 37.0% | $3.0\times10^{-4}$ | 39 | 46 |
| Afatinib no EGF | 0.019±0.011 | Afatinib + EGF | 0.021±0.009 | 6.28% | 0.52 | 53 | 52 |
| Cmpd 2 no EGF | 0.012±0.006 | Cmpd 2 + EGF | 0.013±0.007 | 5.92% | 0.59 | 43 | 50 |

**Supplementary Table 5.**

SPT diffusion results for experiments assessing effects of autophosphorylation and adaptor protein recruitment on EGFR mobility. Measurements in the absence of ligand used QD-conjugated HA tag antibody Fab fragments to label the receptor. Measurements with EGF employed QD-EGF at 200 pM.

| Condition 1 | *D* ($\mu^2 s^{-1}$) | Condition 2 | *D* ($\mu^2 s^{-1}$) | % Change | *P* value | $n_1$ | $n_2$ |
|---|---|---|---|---|---|---|---|
| WT no EGF | 0.028±0.023 | WT +EGF | 0.018±0.007 | 36.4% | $4.9 \times 10^{-6}$ | 124 | 56 |
| ΔC-Tail no EGF | 0.026±0.014 | ΔC-Tail + EGF | 0.023±0.014 | 14.3% | 0.023 | 166 | 118 |
| Y9S no EGF | 0.025±0.012 | Y9S + EGF | 0.018±0.008 | 27.7% | $1.8 \times 10^{-4}$ | 72 | 60 |
| Y10X no EGF | 0.018±0.0066 | Y10X + EGF | 0.015±0.006 | 18.8% | 0.007 | 59 | 68 |
| WT no afat (+ EGF) | 0.027±0.013 | WT + afat (+ EGF) | 0.060±0.013 | 54.5% | $6.7 \times 10^{-21}$ | 46 | 46 |
| KO no afat (+EGF) | 0.052±0.021 | KO + afat (+EGF) | 0.088±0.026 | 40.9% | $1.6 \times 10^{-9}$ | 36 | 42 |

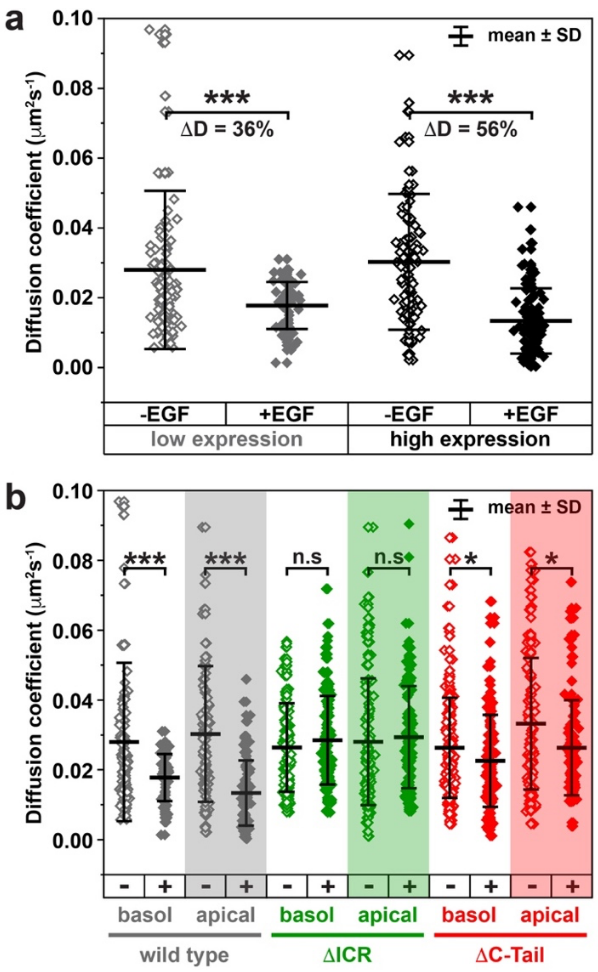


**Supplementary Figure 1**

**Neither expression level nor cellular location affect diffusion coefficient changes.**

**a** SPT studies in CHO cells expressing either low levels (5,000 – 10,000 receptors per cell; gray) or higher levels of wild type EGFR (>100,000 receptors per cell; black). Significant changes in diffusion coefficient are seen for both upon QD-EGF stimulation. $P = 9.2 \times 10^{-6}$ ($n$ = 124 without ligand, 56 with ligand) for low-expressors, and $P = 2.3 \times 10^{-6}$ ($n$ = 98 without ligand, 88 with ligand) for high-expressors.

**b** Receptors were tracked and compared on both the basolateral membrane (using TIRF imaging) and the apical membrane (using wide-field fluorescence imaging). Significant changes in $D$ were seen with QD-EGF for wild type EGFR (gray), with $P = 9.2 \times 10^{-6}$ for basolateral EGFR ($n$ = 124 without ligand, 56 with ligand) and $P = 2.3 \times 10^{-6}$ for apical EGFR ($n$ = 98 without ligand, 88 with ligand) and for ΔC-Tail EGFR (red), with $P$ = 0.023 for basolateral EGFR ($n$ = 166 without ligand, 118 with ligand) and $P$ = 0.0018 for apical EGFR ($n$ = 116 without ligand, 110 with ligand). By contrast, ΔICR EGFR (green) showed no significant change in $D$ with QD-EGF in either location, with $P$ = 0.21 for basolateral EGFR ($n$ = 116 without ligand, 121 with ligand) and $P$ = 0.55 for apical EGFR ($n$ = 113 without ligand, 109 with ligand). Unpaired two-sided Welch's t-tests were used to calculate $P$ values (*$P < 0.05$; **$P < 1 \times 10^{-3}$; ***$P < 1 \times 10^{-6}$), and n.s. indicates no significant difference. Source data are provided as a Source data file.

**a**

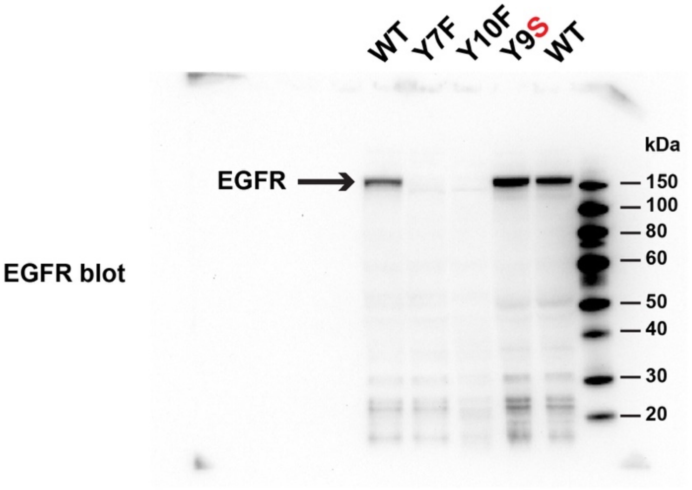


**b**

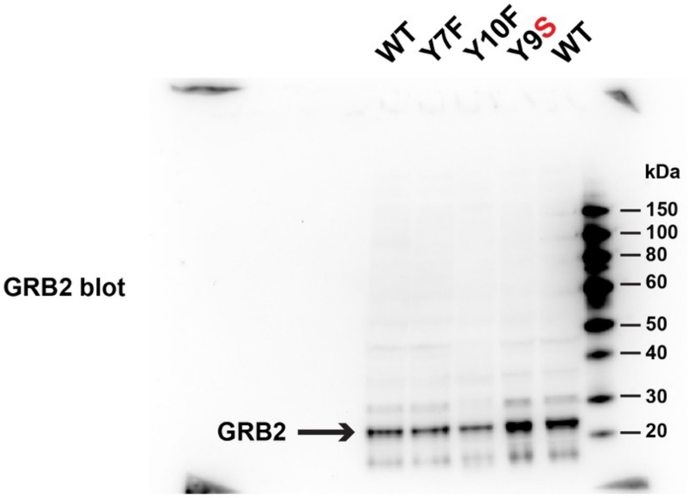


**c**

**Y7F: Y845F/Y992F/Y1045F/Y1068F/Y1148F/Y1173F**
**Y10F:Y845F/Y974F/Y992F/Y1045F/Y1068F/Y1086F/Y1101F/Y1114F/Y1148F/Y1173F**
**Y9S: Y974S/Y992S/Y1045S/Y1068S/Y1086S/Y1101S/Y1114S/Y1148S/Y1173S**

**Supplementary Figure 2**
**Expression of EGFR variants with mutated autophosphorylation sites.**
**a** Uncropped Western blots of whole CHO cell lysates to assess expression of EGFR variants with mutated autophosphorylation sites. The noted variant (or wild type) was expressed transiently in CHO cells, and whole cell lysates subjected to SDS-PAGE using a 4-12% gradient Bis-Tris NuPAGE gel (Thermo Fisher) followed by immunoblotting with antibody AF231 (R&D Systems) as described in Methods. Molecular weight markers are shown at right.
**b** The lysates in **a** were also blotted for GRB2 as loading control (see Methods).
**c** List of mutations included in each EGFR variant analyzed in a, using mature EGFR numbering.

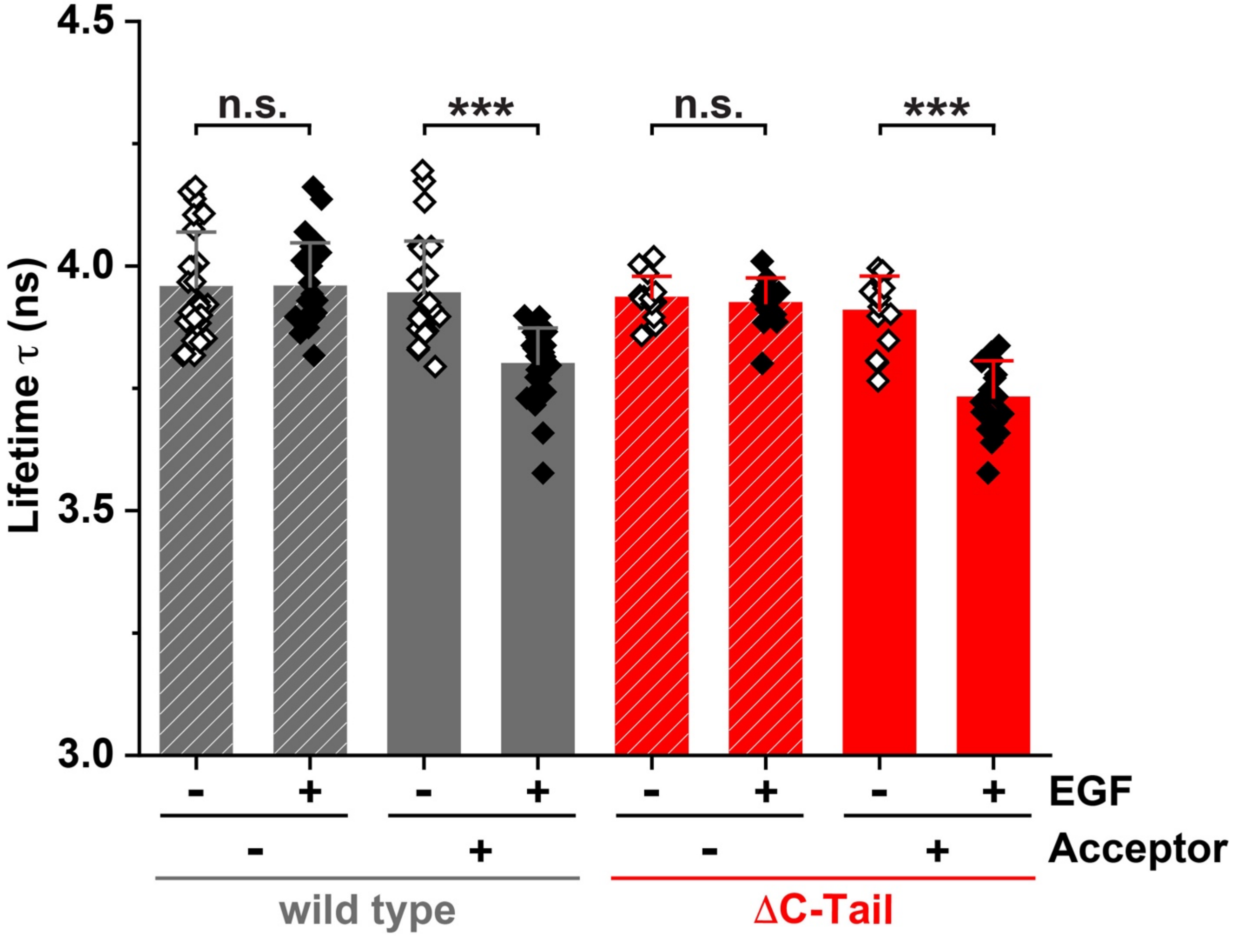


**Supplementary Figure 3**
**FRET-FLIM shows that ΔC-Tail EGFR forms EGF-induced dimers.**
FRET-FLIM imaging analysis of HaloTagged wild type or ΔC-Tail EGFR expressed in CHO cells as described in Methods. Imaging experiments were performed both after labeling with only the donor fluorophore (JF-549; 20 nM) and after labeling with both donor and acceptor (JF-646) fluorophores at equal concentrations (20 nM) – corresponding to '+ Acceptor'. Imaging was performed with (+) and without (-) addition of unlabeled EGF at a saturating concentration (50 nM). Unpaired two-sided Welch's t-tests were used to calculate *P* values (***$P < 1 \times 10^{-6}$), with n.s. indicating no significant difference ($P > 0.05$). Donor fluorescence lifetimes (τ, in ns) were reduced significantly upon EGF addition (when donor and acceptor were both present) for both wild type EGFR ($P = 8.1 \times 10^{-7}$; $n$ = 25 without ligand, 29 with ligand) and ΔC-Tail EGFR ($P = 3.6 \times 10^{-8}$; $n$ = 16 without ligand, 18 with ligand). Source data are provided as a Source data file.

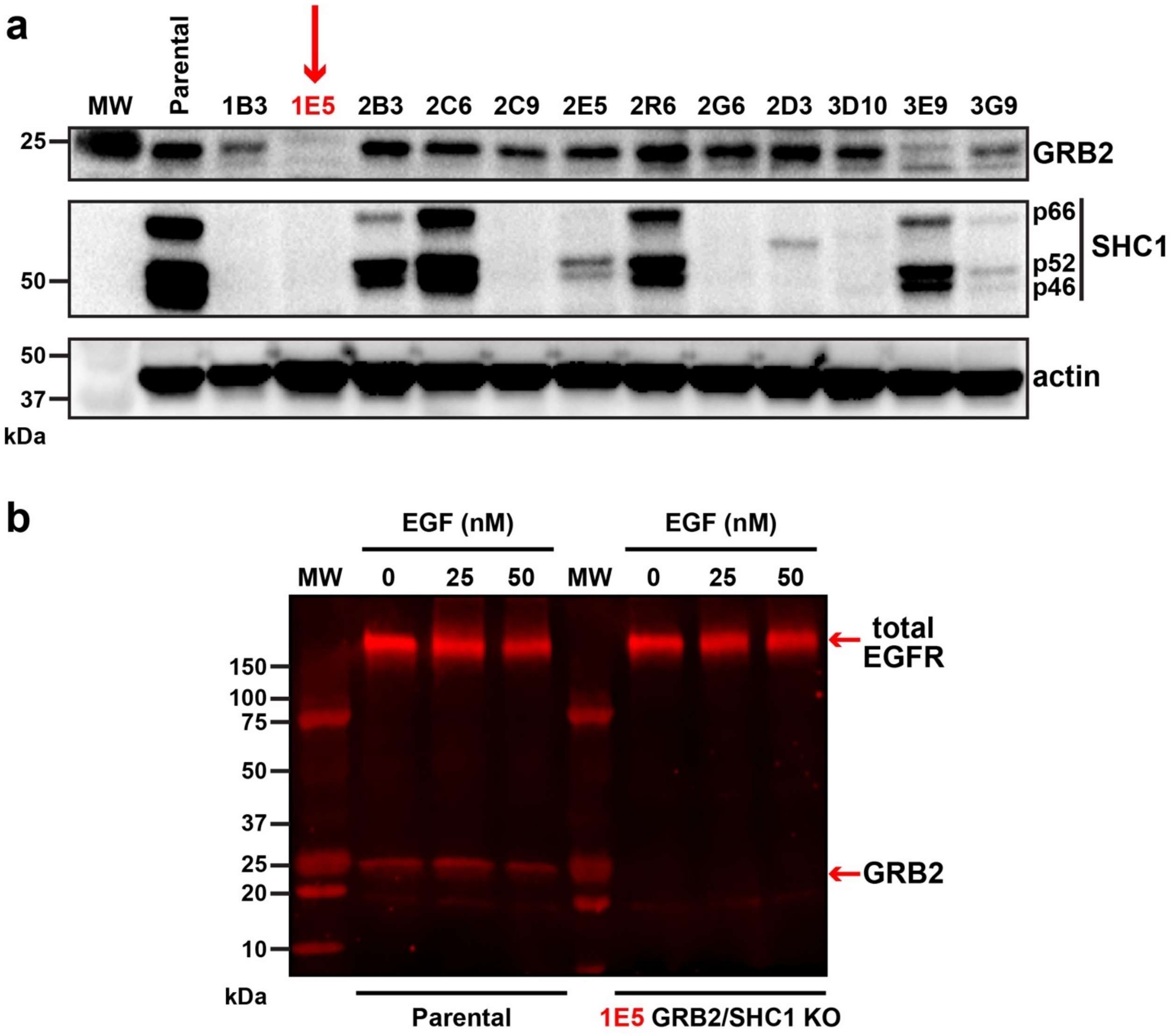


**Supplementary Figure 4**

**Confirmation of CRISPR knock-out of GRB2 and SHC1 in HeLa-CCL-2 cells.**

**a** Whole cell lysates from different selected subclones after CRISPR knock-out of GRB2 and SHC1 (alongside parental cells) were separated by SDS-PAGE on 4-20% acrylamide gradient gels and subjected to immunoblotting with anti-GRB2 (Santa Cruz #SC-255) in the upper panel, anti-SHC1 (BD Biosciences #610878) in the middle panel, and actin (Sigma Aldrich #A1978) as a loading control in the lower panel. As can be seen in the blots, subclone 1E5 lacks detectable expression of both GRB2 and SHC1.

**b** Immunoblot of parental (left) and GRB2/SHC1 1E5 cells for GRB2 (lower) and EGFR (upper) by simultaneous visualization using a LI-COR system, using antibodies described in Methods. Molecular weight markers are shown.

Uncropped gels are provided in the Source Data file.
